# A Chameleonic Macrocyclic Peptide with Drug Delivery Applications

**DOI:** 10.1101/2020.12.16.422786

**Authors:** Colton D. Payne, Bastian Franke, Mark F. Fisher, Fatemeh Hajiaghaalipour, Courtney E. McAleese, Angela Song, Carl Eliasson, Jingjing Zhang, Achala S. Jayasena, Grishma Vadlamani, Richard J. Clark, Rodney F. Minchin, Joshua S. Mylne, K. Johan Rosengren

## Abstract

Head-to-tail cyclized peptides are intriguing natural products with unique properties. The PawS-Derived Peptides (PDPs) are produced from precursors of seed storage albumins in species of the daisy family. Here we report an unusually large PDP with two disulfide bonds, identified from seeds of *Zinnia elegans*. In water, synthetic PDP-23 forms a unique dimeric structure in which two monomers containing two β-hairpins cross-clasp and enclose a hydrophobic core, creating a square prism. This stable dimer can be split and each monomer unfolds to a V-shape in micelles or organic solvents. This chameleonic character is unusual for disulfide-rich peptides and engenders PDP-23 with potential for cell delivery and accessing novel targets. We demonstrated this by conjugating a rhodamine dye to the PDP-23 scaffold, creating a stable, cell-penetrating inhibitor of the P-glycoprotein drug efflux pump.

## Introduction

Macrocyclic and disulfide-rich peptides have generated significant interest for the development of peptide drugs because of their unique stability^1-3^. Naturally occurring macrocyclic peptides ranging in size from 5-74 residues and incorporating up to three disulfide bonds have been discovered in bacteria^4^, fungi^5^, mammals^6^ and plants^7^. One recently discovered family of ribosomally synthesized macrocycles are the PawS-Derived Peptides (PDPs) from the daisy family Asteraceae^8^. The prototypic PDP, Sunflower Trypsin Inhibitor-1 (SFTI-1), was discovered in seeds of the common sunflower *Helianthus annuus*^9^. The sequence for SFTI-1 is within a precursor for a seed storage albumin^10^ called Preproalbumin with SFTI-1 (PawS1). The peptide is excised and head-to-tail cyclized during post-translational albumin processing by asparaginyl endopeptidase^11^.

Based on their distribution, the PDP family is over 18 million years old^12^ and smaller, related peptides pre-date them by another ∼25 million years^13,14^. Like SFTI-1, all PDPs are only found in seeds and are encoded by precursor genes that additionally encode a seed storage albumin. To date, 22 unique PDP sequences have been identified^10,12,15,16^, with 14 confirmed *in planta*. Nineteen PDPs are confirmed or predicted to possess a cyclic backbone, where a proto-N-terminal glycine and a proto-C-terminal aspartate are joined by a transpeptidation reaction performed by asparaginyl endopeptidase. This N-terminal glycine is absolutely conserved amongst all PDPs, as is the C-terminal aspartate for cyclic PDPs. All reported PDPs are ‘stapled’ by a single disulfide bond, which creates a constrained stable structure^8^.

Natural macrocycles have inspired medicinal chemists because of their potential in drug design and engineering applications. Strategies include cyclization of the peptide backbone in otherwise linear peptides with desirable bioactivity to increase stability^17,18^. Alternatively, a bioactive motif can be ‘grafted’ i.e. incorporated into the backbone of stable cyclic peptide scaffolds^18,19^. The grafting concept has been applied to a number of macrocycles, including both SFTI-1 and the larger cyclotides, and used in disease models for cancer^20,21^, multiple sclerosis^22^, or to improve proangiogenic activity^23^. Rather than replacing the entire bioactivity with a new segment, the inherent protease inhibitory activity of SFTI-1 can also be tuned to specifically target different proteases^24,25^. Peptides can also be conjugated to small molecule drugs and act as carriers. Typically, conjugates are focused on overcoming common small molecule issues, such as poor solubility, metabolism and off-target effects due to unfavorable distribution^26^. Peptide-drug conjugates tested in clinical trials include somatostatin and analogues bound to isotopes ^90^Y, ^111^In, and ^177^Lu for targeted radiotherapy^27^, gonadotropin-releasing hormone bound to doxorubicin for the treatment of ovarian and endometrial cancers^28^, as well as angiopep-2 bound to paclitaxel for the treatment of glioblastoma, lung and ovarian cancers^29^. The latter conjugate allows paclitaxel to cross the blood-brain barrier but prevents P-glycoprotein (P-gp) efflux^29^.

Here we describe the identification, chemical synthesis, structural characterization, and a peptide-drug conjugate application of PDP-23, a unique PDP twice the size of SFTI-1 and the first macrocyclic peptide described with two disulfide bonds. PDP-23 was identified by *de novo* transcriptomics in seeds of the common zinnia *(Zinnia elegans)*. Chemical synthesis and NMR spectroscopy revealed PDP-23 self-associates to form a unique symmetrical homodimeric structure. The fold is highly resistant to temperature and enzymatic degradation. However, the quaternary structure disassociates in membrane-mimicking environments, and PDP-23 can penetrate cells. The combination of different loop lengths, multiple turns and a chameleonic structure offer novel design and engineering opportunities to create stable therapeutic leads. This was demonstrated by attaching the small molecule 5/6-carboxy-tetramethyl-rhodamine, a substrate of the multidrug resistance transporter P-gp^30^, to the PDP-23 scaffold, creating a successful, cell-penetrating inhibitor of P-gp.

## Results

### Discovery of PDP-23 by *de novo* transcriptomics

The assembly of whole *de novo* transcriptomes of dry seeds has been shown to be an effective way to discover members of the PDP family^12,31^. A mature PDP sequence can be predicted based on conserved flanking residues that are necessary for processing^10,12,32^. While searching a *Zinnia elegans* transcriptome for the *PawS1* genes that encode PDPs, we found a sequence appearing to encode an unusually large PDP with 28 amino acid residues and the sequence GFCWHHSCVPSGTCADFPWPLGHQCFPD. A detailed analysis of the RNA-seq reads confirmed the sequence with an average depth of coverage of 696-fold at each nucleotide position, giving high confidence (**Supplementary Fig. 1**). The theoretical monoisotopic mass of the linear reduced peptide is 3130.29 Da. However, for all reported PDPs where the residue at the C-terminal processing site is an Asp, it is joined head-to-tail with Gly1 in a cleavage-coupled intramolecular transpeptidation reaction^11^, and the four cysteine residues are expected to form two disulfide bonds. Thus, the expected monoisotopic mass for a processed cyclic and oxidized peptide is 3108.24 Da. Analyzing seed extracts by LC-MS, we detected a series of [M+3H]^3+^ ions whose mass matched the prediction (**Fig. 1**), indicating this peptide does indeed exist. It was named PDP-23, in keeping with previous nomenclature^12,15^.

**Figure 1:**
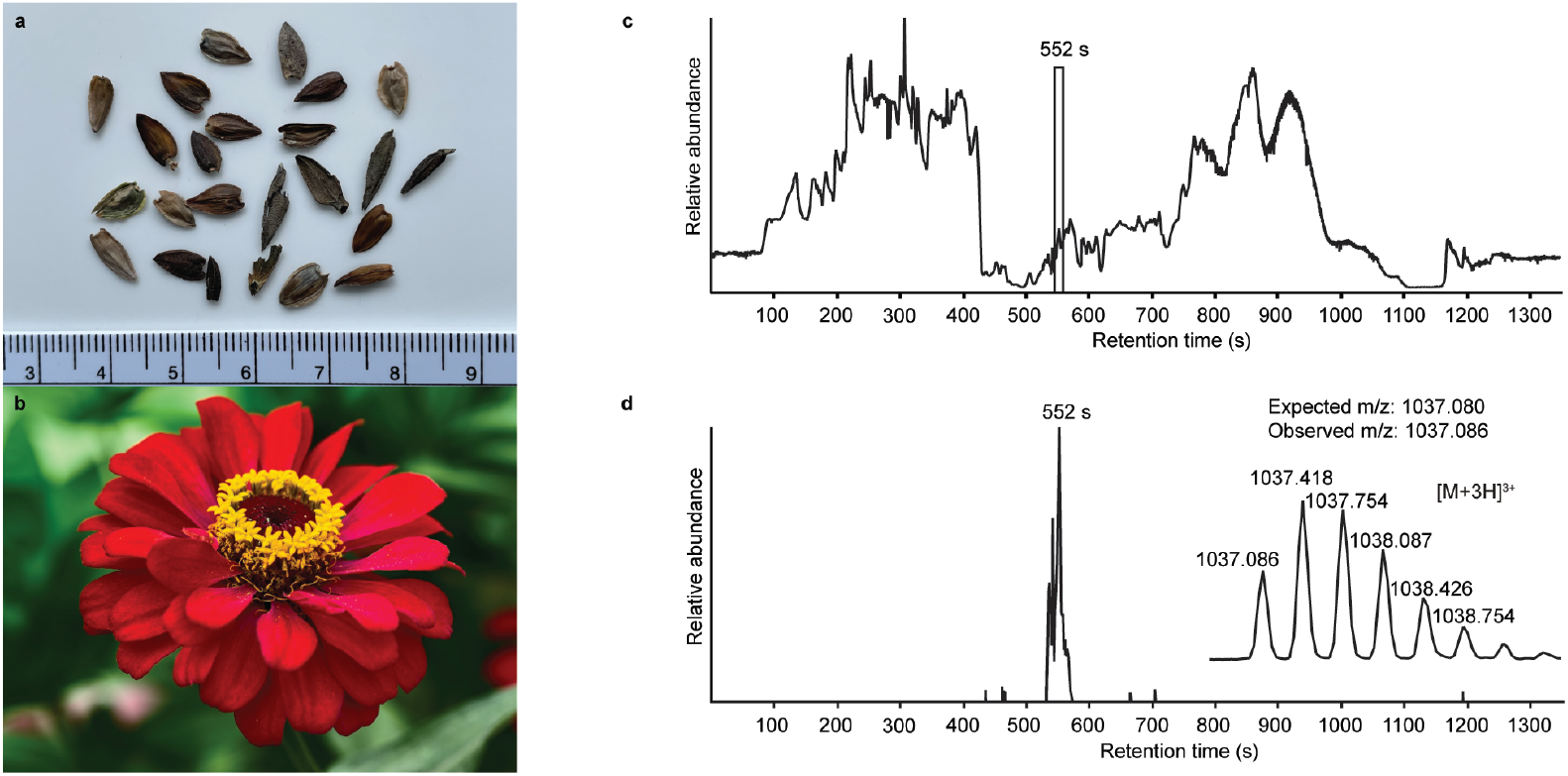
LC-MS data confirming the presence of PDP-23 in a seed extract of *Zinnia elegans*. **(a)** *Zinnia elegans* seed, ruler placed for scale. **(b)** Image of a *Zinnia elegans* flower. **(c)** Total ion chromatogram of the seed extract marked with the retention time of PDP-23, confirming its presence at the protein level. **(d)** Extracted ion chromatogram (XIC) for PDP-23 showing the [M+3H]^3+^ (*m/z*) state of the observed mass-to-charge ratio and its isotopic peak envelope.

### Extraction of PDP-23 and LC-MS/MS sequencing

To confirm the sequence of PDP-23, the peptide was extracted from seeds, partially purified using RP-HPLC and the monoisotopic mass confirmed by MALDI-TOF. The PDP-23 enriched extract was reduced, alkylated and digested with chymotrypsin, and the cleaved mixture was subjected to sequencing by LC-MS/MS. The resulting spectra were searched against a custom-built peptide database and three peptide fragments (HHSCVPSGTCADFPWPL, GHQCFPDGFCW, HHSCVPSGTCADFPWPLGHQCFPDGFCW) that cover the entire sequence of PDP-23 were identified, including the macrocyclization junction between Asp28 and Gly1. Each MS/MS spectrum was assigned b-and y-ion series (**Supplementary Fig. 2**). As mass spectrometry cannot readily distinguish between Gln/Lys, to confirm the presence of Gln24 we reduced, alkylated and digested the PDP-23 enriched extract with trypsin, but no tryptic fragments were identified, consistent with the PDP-23 sequence not containing any Lys or Arg residues. MS also does not distinguish between Leu/Ile residues, but the transcriptomic raw reads strongly support a Leu residue at position 21 (**Supplementary Fig. 1**) and successful cleavage by chymotrypsin to generate the fragment HHSCVPSGTCADFPWPL also supports this assertion.

### Peptide synthesis and purification of PDP-23 disulfide conformers

PDP-23 was found to be of very low abundance in seeds. Thus, to get material to further characterize this peptide we turned to chemical synthesis. With two disulfide bonds, three different disulfide connectivities are theoretically possible (**Fig. 2A**). Each of the three possible PDP-23 conformers was synthesized separately using solid phase Fmoc chemistry, cyclized in solution, and oxidized by regioselective disulfide bond formation. Each conformer was purified by RP-HPLC and the monoisotopic mass confirmed by MALDI-TOF MS (**Fig. 2B-C**). Conformers I and II showed a monodispersed peak shape, whereas conformer III eluted with a tailing shoulder. To confirm that the three synthesized conformers possess the same purity, sequence and cyclic backbone, they were reduced, alkylated and analyzed by RP-HPLC. The analytical RP-HPLC trace showed identical elution at 40.55 min for all three reduced and alkylated conformers.

**Figure 2:**
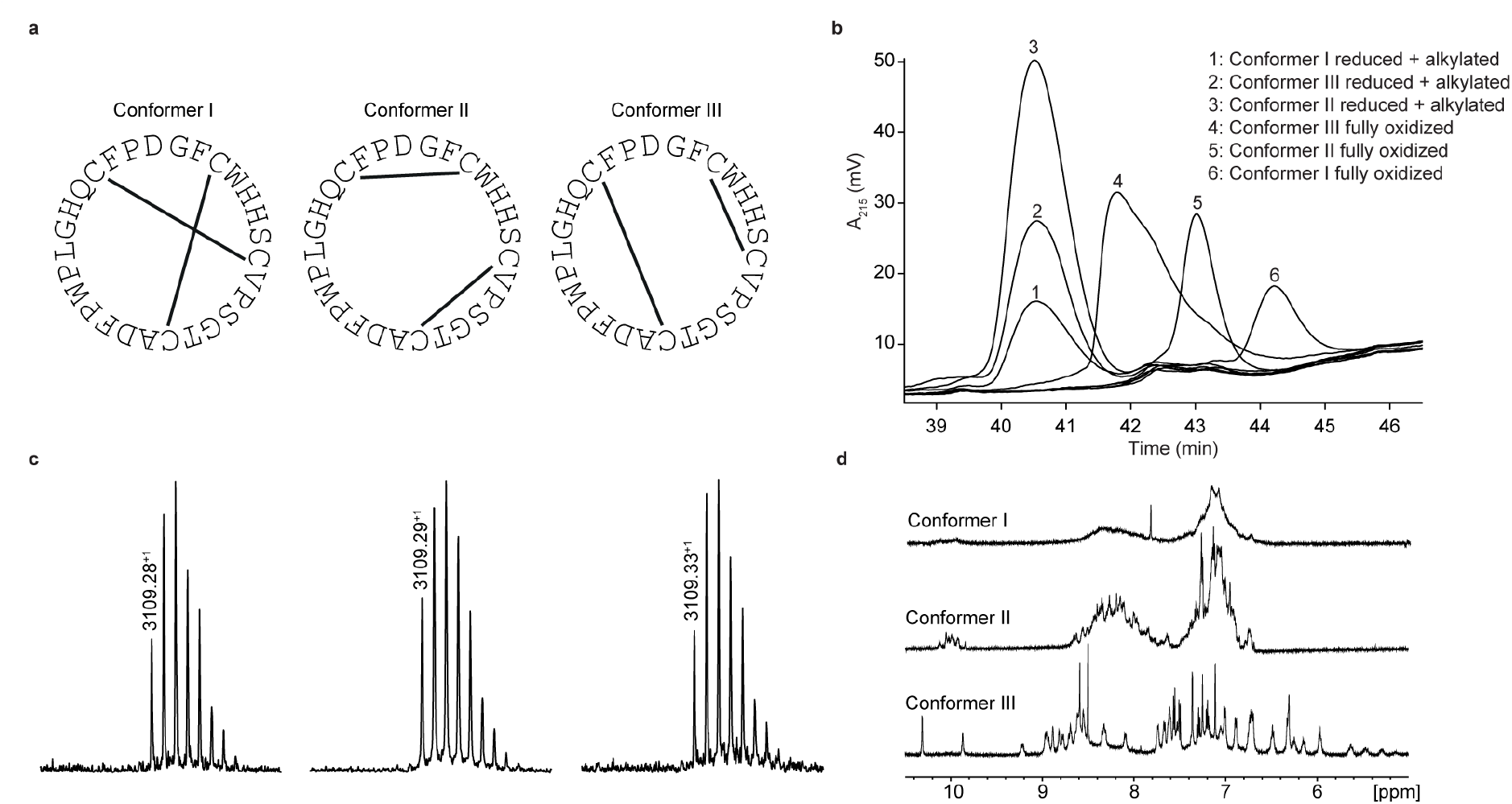
Synthesized conformers of the PDP-23 peptide. **(a)** Cysteine connectivity of the three synthesized conformers with corresponding sequence. **(b)** Analytical RP-HPLC trace of synthesized conformers in reduced and alkylated and fully oxidized state, respectively. The conformers eluted in the following order: conformer III (41.78 min), conformer II (43.02 min) and conformer I (44.22 min) **(c)** MALDI-TOF mass spectra showing a well-distributed experimental isotopic peak envelope with an observed monoisotopic mass-to-charge ratio of 3109.28^+^ for conformer I, 3109.29^+^ for conformer II and 3109.33^+^ for conformer III. **(d)** One-dimensional NMR spectra of the three fully oxidized conformers.

To confirm which of the synthetic versions represents the native variant we compared the elution times of the oxidized peptides with the PDP-23 enriched seed extract using analytical RP-HPLC. The earliest eluting conformer, conformer III, was identified as the closest match to our enriched extract (**Supplementary Fig. 3A**), however it was not possible to conduct co-elution studies as PDP-23 remained only a minor component of the extract and could not be purified from more abundant components. MALDI-TOF analysis highlighted the low abundance of PDP-23 within the enriched extract at a mass-to-charge ratio of 3109.35^+^, along with a number of more abundant masses (**Supplementary Fig. 3B**).

### NMR spectroscopy studies of PDP-23

Solution NMR spectroscopy was used to study all three synthetic conformers to gain further insight into the likely native conformation. Conformer III showed excellent dispersion in the 1D ^1^H NMR spectrum, indicating a folded, highly structured peptide, whereas conformers I and II showed poor dispersion consistent with disorder (**Fig. 2D**). In addition, conformer I showed very broad lines consistent with aggregation and conformer II generated multiple signals from the Trp indole protons suggesting multiple conformational states. Clearly the disulfide array of conformer III is structurally favored. All PDPs that have been isolated from seeds have been shown to be highly structured, whereas misprocessed forms tend to be degraded^11^, which strongly supports conformer III being the native form. Extensive NMR data were recorded for conformer III and assigned manually using sequential assignment strategies. The high-quality datasets allowed for the complete assignment of backbone and sidechain resonance using TOCSY and NOESY spectra (**Supplementary Fig. 4**)^33^. HSQC data were recorded at natural abundance, allowing assignments of the ^13^C and ^15^N backbone and side chain resonances based on ^1^H assignments. PDP-23 contains four proline residues with Pro20 confirmed to be in the *cis* configuration based on NOE patterns and ^13^C chemical shifts, whilst Pro10, Pro18 and Pro27 all adopt a *trans* conformation. No additional spin systems suggesting conformational inhomogeneity were identified.

To determine the 3D structure of PDP-23, structural restraints were derived from the NMR data. These included inter-proton distances based on NOEs, dihedral angles based on chemical shifts, and hydrogen bond donors based on temperature coefficients. Intriguingly, after considerable efforts, it was realized that a 3D structure consistent with all data could not be computationally generated. A number of unambiguous NOEs were unable to be satisfied by the structures generated. For example, NOEs were observed from the H*β*s of Trp4, the H*β*s of Ser7, and the Hγ protons of Val9 to the Hδ methyl protons of Leu21. However, the distances between these protons all fell well outside the possible range of the NOE (∼5 Å) in the calculated models. The only possible explanation was that these are intermolecular NOEs, and due to the lack of any resonance duplication PDP-23 must exist in solution as a symmetric multimer.

### Structure determination of PDP-23 in monomeric and dimeric forms

In an attempt to separate the multimeric PDP-23 into individual monomers we explored adding organic solvent. Recording NMR spectra of PDP-23 in a solution of 80:20 H_2_O/CD_3_CN revealed a significant change in dispersion and peak line widths (**Supplementary Fig. 5**). The CD_3_CN data showed narrower lines, while overall peak dispersion was decreased, in particular in the aromatic region, indicating a change in the structure. Rerecording and reassigning all 2D NMR revealed significant changes in the chemical shifts of numerous protons, and also in the NOE patterns observed. Critically, key NOEs identified as intermolecular were no longer present and residues involved experienced chemical shift deviations > 0.15 ppm for numerous protons. However, the secondary Hα shifts, which are sensitive indicators of secondary structure, are remarkably similar between PDP-23 in water and in 80:20 H_2_O/CD_3_CN (**Supplementary Fig. 6**). Thus, as a whole, the backbone and secondary structure of PDP-23 remains largely the same.

Structural restraints were derived from the NMR data and 3D structures were calculated using simulated annealing. In contrast to the calculations performed using the aqueous data, this resulted in structures fully consistent with all observed NOEs. The ensemble of the 20 best models chosen to represent the monomeric structure of PDP-23 in 80:20 H_2_O/CD_3_CN is shown in **Fig. 3A**. PDP-23 comprises two double stranded anti-parallel β*-*sheets, instead of the one β*-*sheet typical of the smaller members of the PDP family. A single disulfide bond bridges each β-sheet and the backbone is stabilized by a large number of hydrogen bonds (**Fig. 3D**). The structure is V-shaped with polar sidechains being surface exposed, whereas the hydrophobic residues Phe2, Trp4, Val9, Phe17, Leu21, and Phe26 create a hydrophobic core sandwiched between the sheets on the inside of the ‘V’. In addition to the two anti-parallel β*-*sheets, several well-defined turns were identified: a type I’ turn comprising residues Trp4-Ser7, a type II turn comprising residues Pro10-Thr13, a type VIa1 turn comprising residues Pro18-Leu21 and a type I turn comprising residues Phe26-Gly1. Three of the turns feature proline residues, which may be key features for creating bends in the backbone.

**Figure 3:**
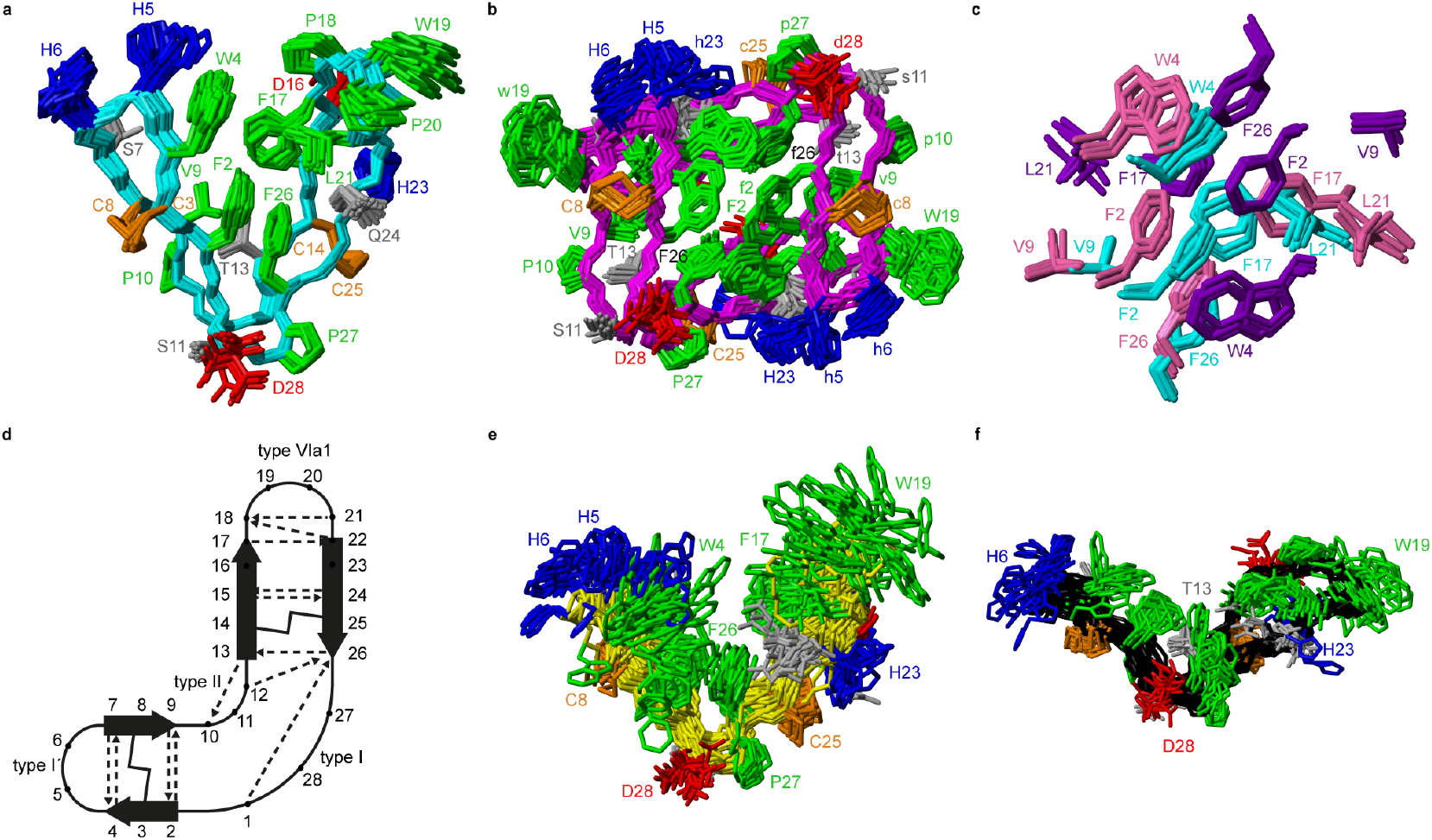
Three-dimensional structure of PDP-23 in different environments. **(a)** Superposition of the 20 best structures of PDP-23 in 80:20 H_2_O/CD_3_CN. The cyclic backbone is highlighted in cyan, disulfide bonds in orange, hydrophobic residues in green, basic residues in blue, acidic residues in red, and polar residues in grey. Residues are labelled by one letter code in colors representative of residue type. **(b)** Superposition of the 20 best structures of the symmetrical homodimer fold of PDP-23 in 90:10 H_2_O/D_2_O. The same nomenclature is used as in panel **(a)**, barring the cyclic backbone, which is displayed in magenta. Upper and lower case lettering are used to distinguish between the two PDP-23 molecules. (**c**) Superposition of the hydrophobic core regions of PDP-23 in 90:10 (H_2_O/D_2_O) and 80:20 (H_2_O/CD_3_CN). PDP-23 in H_2_O/D_2_O is shown in cyan with PDP-23 in H_2_O/CD_3_CN being shown in magenta and purple to delineate the two monomers. All residues are labeled in color to assist with orientation **(d)** Schematic representation of monomeric PDP-23 showing structural motifs including the two anti-parallel β-sheets, disulfide bonds, turns and hydrogen bond network. **(e)** Superposition of the 20 best structures of PDP-23 when exposed to SDS micelles. The same nomenclature is used as in panel **(a)**, barring the cyclic backbone, which is displayed in yellow. **(f)** Superposition of the 20 best structures of PDP-23 when exposed to DPC micelles. The same nomenclature is used as in panel **(a)**, barring the cyclic backbone, which is displayed in black.

Returning to the NMR data recorded in H_2_O, we used the structural restraints derived to instead calculate a dimeric structure, enforcing symmetry whilst still allowing for ambiguous NOEs to be assigned as intra-or intermolecular contacts. The generated set of models of a symmetrical dimer were able to satisfy all of the observed NOEs. A large number of NOEs including the aforementioned contacts between the Trp4, Ser7, Val9 and Leu21 were confirmed to result from intermolecular cross relaxation at the interface between two PDP-23 molecules. The structures were again refined and the best 20 models chosen to represent the dimeric structure of PDP-23 in H_2_O (**Fig. 3B**). The structural statistics of PDP-23 as both a monomer and a dimer highlight the high quality of the structures and the excellent agreement of the observed data in both environments (**Supplementary Table 1**). Both structures are well defined with backbone RMSDs of 0.43 and 0.72 Å for the monomer and dimer, respectively. Comparing the dimer and monomer structures, as indicated by the secondary shifts, the backbone structure is very similar and identical hydrogen bonds are present in both forms. The differences are related to the V-shape which opens up in the dimer, increasing the distance between the loops from ∼13 to ∼21 Å. This allows the creation of a larger combined hydrophobic core arranging the two ‘Vs’ into a square prism shape (**Fig. 3B**). The cores of both the monomer and the dimer contain the same residues. However, the larger number of aromatic residues in the dimer’s core and packing variations explain the differences in chemical shifts of many of these protons.

### Structure determination of PDP-23 in micelles

Given that PDP-23 can change structure based on conditions and its tertiary structure is not covalently restrained by the disulfide bonds, we speculated that the large number of hydrophobic amino acids would allow it to interact with membranes. To mimic this situation, we prepared NMR samples containing sodium dodecyl sulfate (SDS) or dodecylphosphocholine (DPC) micelles. Exposure of PDP-23 to SDS micelles resulted in significant spectral changes. Line broadening occurred, consistent with an increase in correlation time as a result of binding to the micelles, and the signal dispersion was reduced. Comparing secondary chemical shifts revealed that the β-sheet structure is largely retained, with similar patterns of secondary Hα chemical shifts (**Supplementary Fig. 6**). Some sidechain resonances, however, drastically changed chemical shift (**Supplementary Fig. 7, Supplementary Table 2**), and many sidechain-to-sidechain NOEs, particularly involving aromatic residues, disappeared. Importantly, no intermolecular NOEs are present. Thus, we conclude that, rather than forming a dimer, the hydrophobic face of PDP-23 is buried in the SDS micelles. Structures were calculated, and although less defined than in solution they indicated that the structure is largely maintained in the presence of SDS micelles, but it is more open, similar to the structure seen in the dimeric form (**Fig. 3E**). This aligns with the 3-4 nm size of an SDS micelle^34^, which would allow for a PDP-23 molecule to expose its hydrophobic core to the micelle interior by slightly opening the V-shape, but retaining the polar groups in solution or around the polar lipid headgroups.

A similar trend occurred when exposing PDP-23 to DPC micelles, which are larger than SDS micelles at ∼6.4-7.2 nm in size^35^. Again, line broadening occurred, in this case for numerous resonances, which created issues with separating spin systems from one another as well as from the noise. Many chemical shift changes in the hinge region and of the aromatic resonances (**Supplementary Fig. 7, Supplementary Table 2**), as well as disappearance of NOEs, including all intermolecular NOEs, were similar to what was observed in the SDS data. The patterns of secondary Hα chemical shifts were again retained (**Supplementary Fig. 6**) and structure calculations using the DPC data indicate that the secondary structure is retained, while the peptide now presents as an almost flat surface, exposing all hydrophobic residues (**Fig. 3F**). The distance between the two loops has extended to ∼32 Å. To analyze differences in hydrogen bonding, amide temperature coefficients were determined under all conditions. These confirmed that the hydrogen bonding network within the sheets is retained in all structures. In contrast the hydrogen bonds stabilizing the turns at the bottom of the ‘V’ from the amide protons of Thr13 and Gly1 to the carbonyls of Pro10 and Phe26, respectively, are lost in the micelle structures (**Supplementary Fig. 8**). This is consistent with the opening of the hinge.

### Thermal and enzymatic stability of PDP-23

As stability is a key feature among macrocyclic peptides we assessed the thermal stability of PDP-23 in H_2_O using NMR spectroscopy. 1D ^1^H spectra were recorded at temperature intervals from 298 K to 363 K (**Supplementary Fig. 9**). PDP-23 was highly stable at elevated temperatures with the downfield H*α* protons characteristic of the *β*-sheet being observed at temperatures up to 363 K. The chemical shift separation of the indole protons of the two tryptophan residues was also retained, however chemical shift changes in the aromatic region are consistent with changes in the hydrophobic core. The symmetrical homodimer quaternary fold was confirmed at temperatures up to 318 K using NOESY data. However, increasing the temperature further to 328 K causes the key dimer NOEs to disappear, indicating a likely dissociation of the dimer. After a decrease in temperature back to 298 K the 1D ^1^H NMR spectrum was found to be identical to the spectrum recorded prior to heating, showing a complete reversibility of any unfolding or destabilization of the dimer.

Given the potential of cyclic peptides as drugs, we wanted to evaluate the stability of PDP-23 in biological settings. We exposed the peptide to serum as well as simulated gastric and intestinal fluids. PDP-23 was found to be highly stable with more than 50% remaining intact after 24 h incubation in either serum or gastric fluid and the half-life in simulated intestinal fluid being >2 h (**Fig. 4A-C**).

**Figure 4:**
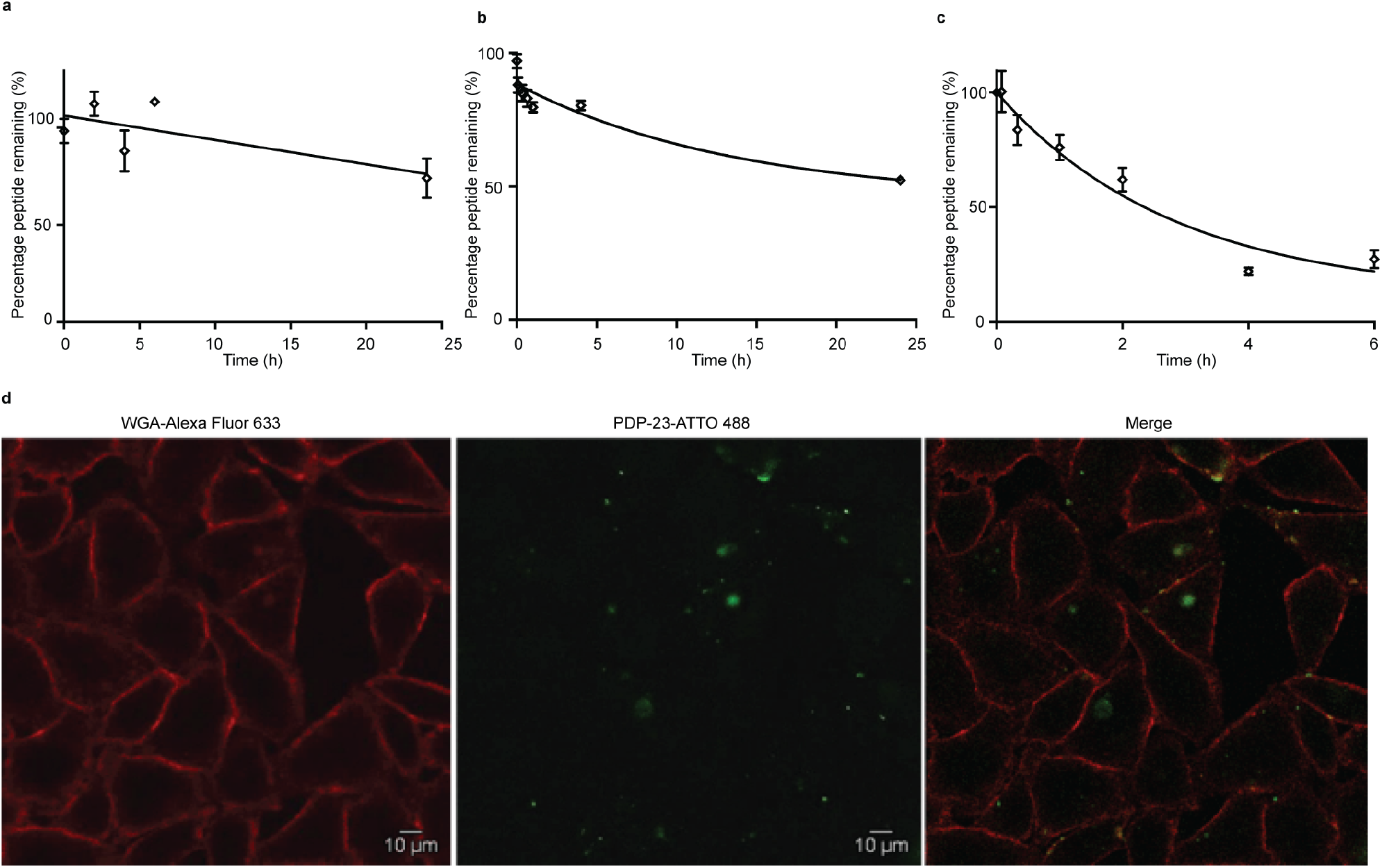
Stability and internalization of PDP-23. Non-linear regression fit of the decay of PDP-23 over time when exposed to **(a)** human serum, **(b)** simulated gastric fluid and **(c)** simulated intestinal fluid. All data are means ± SEM, n = 3. **(d)** Internalization of PDP-23 in HeLa cells. PDP-23 labelled with ATTO 488 (2 µM concentration) was incubated with the cells for 2 h at 37 °C. The plasma membrane was labelled with WGA-Alexa 633 as seen in red. The labelled PDP-23 is visible in green. The magnification used was 63X.

### PDP-23 is non-toxic and can penetrate cells

Given PDP-23’s ability to insert into micelles we considered it may interfere with membrane integrity and be toxic to cells. To investigate the cytotoxicity of PDP-23 an MTT assay was conducted using HeLa cells. Cells were exposed to concentrations of PDP-23 ranging from 0.5 to 32 µM for 48 h at 37 °C, but no significant cytotoxic activity was observed (**Supplementary Fig. 10**). Fluorescence microscopy was used to investigate the ability of PDP-23 to internalize. PDP-23 was synthesized with Asp28 replaced by a Lys (PDP-23 D28K) to allow labeling with ATTO 488 and incubated with HeLa cells. The distinct green fluorescence of ATTO 488 labeled PDP-23 was found internalized in green puncta within HeLa cells, suggesting endosomal uptake. Furthermore, diffuse green coloring of the cytosol suggested the peptide is able to reach the cytoplasm (**Fig. 4D**).

### PDP-23 as a scaffold for therapeutic drugs

PDP-23, due to its stability, lack of toxicity and ability to penetrate cells, was thought to be an excellent candidate as a scaffold for drug development. A substrate of the multidrug exporter P-glycoprotein (P-gp), the dye 5/6-carboxy-tetramethyl-rhodamine, was conjugated to PDP-23 D28K. We speculated that the conjugate would be able to enter cells and while the rhodamine would interact with P-gp, the large peptide would prevent efflux. Thus, we tested PDP-23-rhodamine as an inhibitor of P-gp by exposing it in combination with daunorubicin, a well-established chemotherapeutic and substrate of P-gp^36^, to the cancer cell lines, KB-3-1 and its derivative KB-V-1. The KB-V-1 cells overexpress P-gp whilst the P-gp expression in KB-3-1 cells was undetectable by Western blot. (**Fig. 5A**). Consequently, the KB-V-1 cells displayed significantly decreased sensitivity to daunorubicin compared to the sensitive KB-3-1 cells, due to rapid export via P-gp^36^ (**Fig. 5B)**. Addition of PDP-23-rhodamine in conjunction with daunorubicin, significantly (P ≤0.002) increased cell death in the KB-V-1 cells in a dose-dependent manner, demonstrating the restoration of sensitivity of the KB-V-1 cells to daunorubicin (**Fig. 5C**). However, PDP-23-rhodamine did not significantly increase cell death in the KB-3-1 cells, indicating the selective effect of the conjugate on the P-gp overexpressing KB-V-1 cells. These results are consistent with what was observed for the classical P-gp inhibitor verapamil^37^.

**Figure 5:**
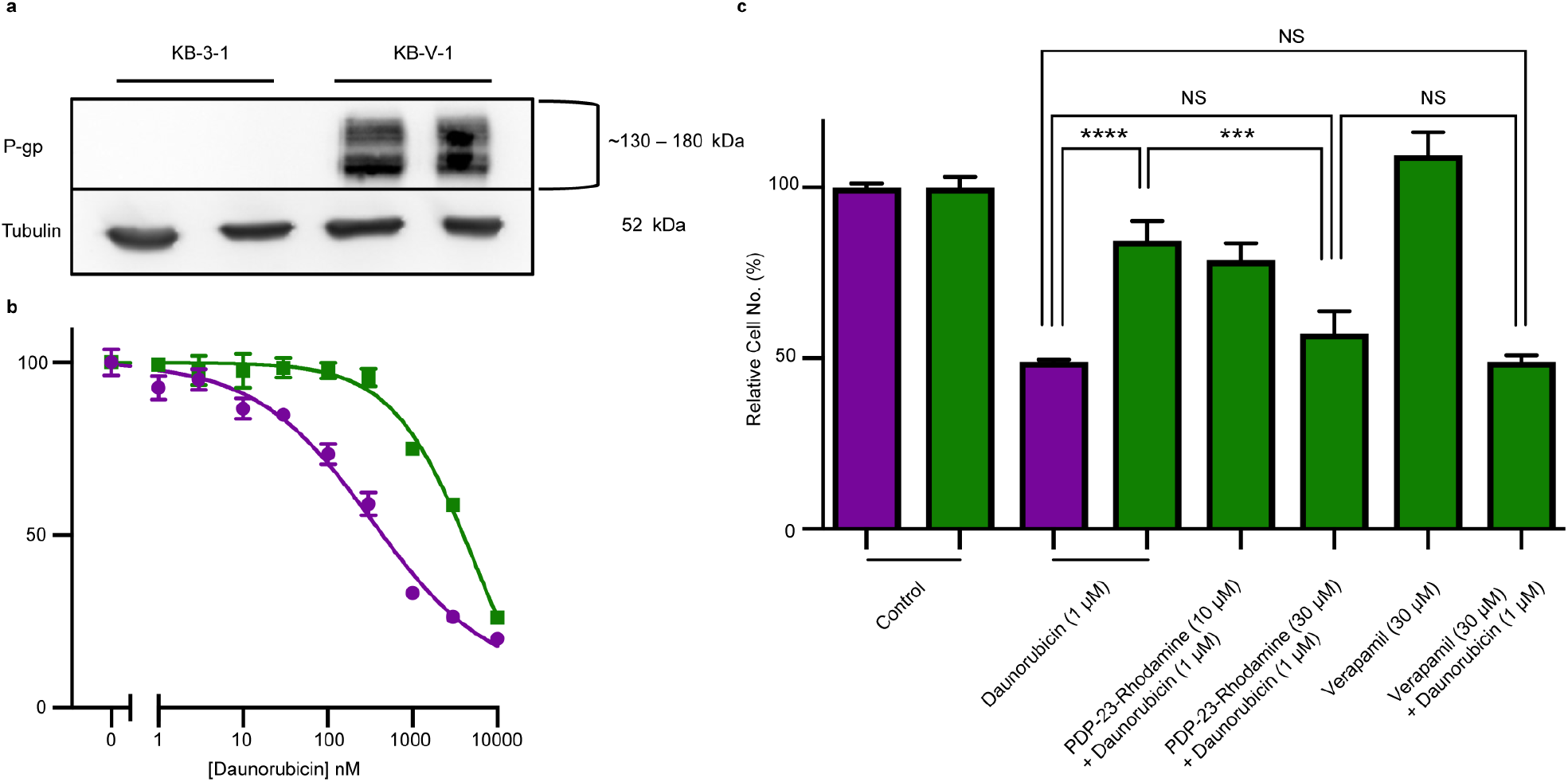
Reversal of daunorubicin resistance when in combination with PDP-23-rhodamine conjugate in P-gp-overexpressing cells. **(a)** Western blot of KB-3-1 and KB-V-1 cells confirming P-gp overexpression in the KB-V-1 cell line. **(b)** Dose-response curve for daunorubicin with KB-3-1 and KB-V-1 cell lines. Cells were treated for 24 h with a range of nine concentrations of the drug. Cell populations determined via quantification of DNA with CyQuant Cell Proliferation Assay Kit and normalized to the untreated control. All data are means ± SEM, n = 4. **(c)** Comparison of toxicity of a single concentration of daunorubicin (1 µM) in sensitive (KB-3-1, 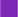) and resistant (KB-V-1, 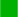) cells, with and without the presence of the PDP-23-rhodamine conjugate. Verapamil (30 µM), a P-gp subtrate, is included as control for P-gp inhibition. Cell populations were determined via quantification of DNA with the CyQuant Cell Proliferation Assay Kit and normalized to an untreated control. All data are mean ± SEM, n = 4; significance determined by two-way ANOVA (**** p<0.0001), (*** p<0.001).

## Discussion

In this study, PDP-23 was identified by transcriptomics and confirmed in *Zinnia elegans* by MS/MS sequencing and MALDI-TOF analysis. At 28 residues PDP-23 is twice the size of typical PDPs and has double the number of cysteine residues. The presence of multiple disulfide bonds presents a challenge when describing novel isolated peptides, as confirming the disulfide connectivity is not trivial. Even when substantial amounts of material are available, allowing large-scale extraction and purification, identification of disulfide bonds may require detailed NMR spectroscopy or chemical approaches^38^. Despite there being sufficient PDP-23 for MS/MS sequencing, the overall abundance of PDP-23 is low and severe chromatographic overlap with other peptidic compounds prevented isolation of a pure native sample. Instead, the three possible disulfide conformers were synthesized by solid phase peptide synthesis and studied by chromatography and NMR spectroscopy. Conformer III was the earliest eluting variant and showed excellent peak dispersion in the 1D ^1^H NMR spectrum, consistent with a well-structured peptide. In contrast the NMR data for conformers I and II showed poor dispersion, conformational inhomogeneity and aggregation. Comparing the RP-HPLC elution times of the different synthetic conformers with the native peptide was a challenge without a pure native sample. The elution time of conformer III showed the best match with the elution time of the enriched PDP-23 extract, and this together with its ordered structure strongly suggests that it represents the native form. The XIC peak for PDP-23 (**Fig. 1**) is unusually broad and does not show a uniform peak, which likely reflects the disassociation of the symmetrical homodimer during elution with acetonitrile, similar to the elution profile with a tailing shoulder observed for synthetic PDP-23 conformer III.

The structural investigation of PDP-23 uncovered different folds of the peptide depending on the environment. In an environment with organic co-solvent, PDP-23 adopts a monomeric structure consisting of an elongated cyclic oval shape with two β-hairpins. A single disulfide bond, supported by an extensive hydrogen bond network, bridges each β-sheet. These loops then enclose a hydrophobic core to form a tertiary V-shaped structure (**Fig. 3A**). In an aqueous environment, PDP-23 forms a symmetrical homodimer through the self-association of two PDP-23 molecules by opening the ‘V’ and interlocking the hydrophobic cores. The quaternary structure of the symmetrical homodimer is roughly a square prism with the hinge of each monomer forming one of two planar vertices and the turn at the top of each loop create a stacked vertex (**Fig. 3B**). The PDP-23 dimer is able to disassociate and interact with micelles by exposing the hydrophobic core of each monomer, while maintaining the β-sheets of each loop. The amount of separation required of the two loops to expose the hydrophobic core of the PDP-23 monomer to a micelle appears to be dependent on the size of the micelle. This is illustrated by the difference in the structural fold observed between PDP-23 when exposed to either SDS or DPC micelles. The dimer also dissociates into individual monomers at increased temperature (>50 °C), but again retains a high degree of structure and folds back into a dimer when the temperature is lowered even after heating to 90 °C. The adaptability of the structure indicates an innate ability of PDP-23 to conform and stabilize its molecular structure depending on the environment it is exposed to.

PDP-23 represents a new structural class of peptides. Previously described PDPs that are distributed throughout the sunflower family are mostly head-to-tail cyclic peptides that are bridged by a single disulfide bond, including the prototypic member SFTI-1^10,12,15,16^. From a structural perspective PDP-23 has more in common with the much larger cyclotides that are found scattered throughout the plant kingdom (**Fig. 6**)^39^. These are also head-to-tail cyclized peptides that are rich in β-sheet structure and tight turns. However, the cyclotides have six conserved cysteines adopting a knotted arrangement of three disulfide bonds^39^. Unlike PDP-23, in the cyclotides the disulfide bonds are buried in the core leaving all hydrophobic residues on the outside. Due to its cysteine connectivity (Cys^I^-Cys^II^ and Cys^III^-Cys^IV^), PDP-23 can be considered as adopting a disulfide-laddered structure. A disulfide-ladder motif is the signature of *θ*-defensins, which are cyclic peptides originally discovered in the leukocytes of rhesus monkeys^6^. The *θ*-defensins comprise only a single β-sheet that is stabilized by three disulfide bonds and are straight in contrast to the V-shaped PDP-23. Intriguingly PDP-23 shares some structural details with both *θ*-defensins and cyclotides. The Trp19-Pro20 turn of PDP-23 is identical to a turn seen in kalata B1, which due to the *cis* peptide bond has been used to define the Möbius cyclotide sub-class. At the other end of PDP-23 the His5-His6 type 1*′* turn and its stabilizing disulfide is conformationally akin to the *θ*-defensin RTD-1 (**Fig. 6**). Whilst disulfide-laddered peptides are rare, cysteine knotted counterparts are not only found in plants but are quite common in nature and are present in various spider, cone snail and plant peptides^40,41^. A degree of self-association has been reported for both cyclotides and *θ*-defensins under NMR conditions but neither involves structural changes or leads to stable defined multimeric forms^42,43^. Importantly, we believe PDP-23 is unique among disulfide-rich peptides in that the disulfides do not directly cross-brace and lock in the tertiary fold, only the secondary structure. The folding around a hydrophobic core is more reminiscent of larger proteins.

**Figure 6:**
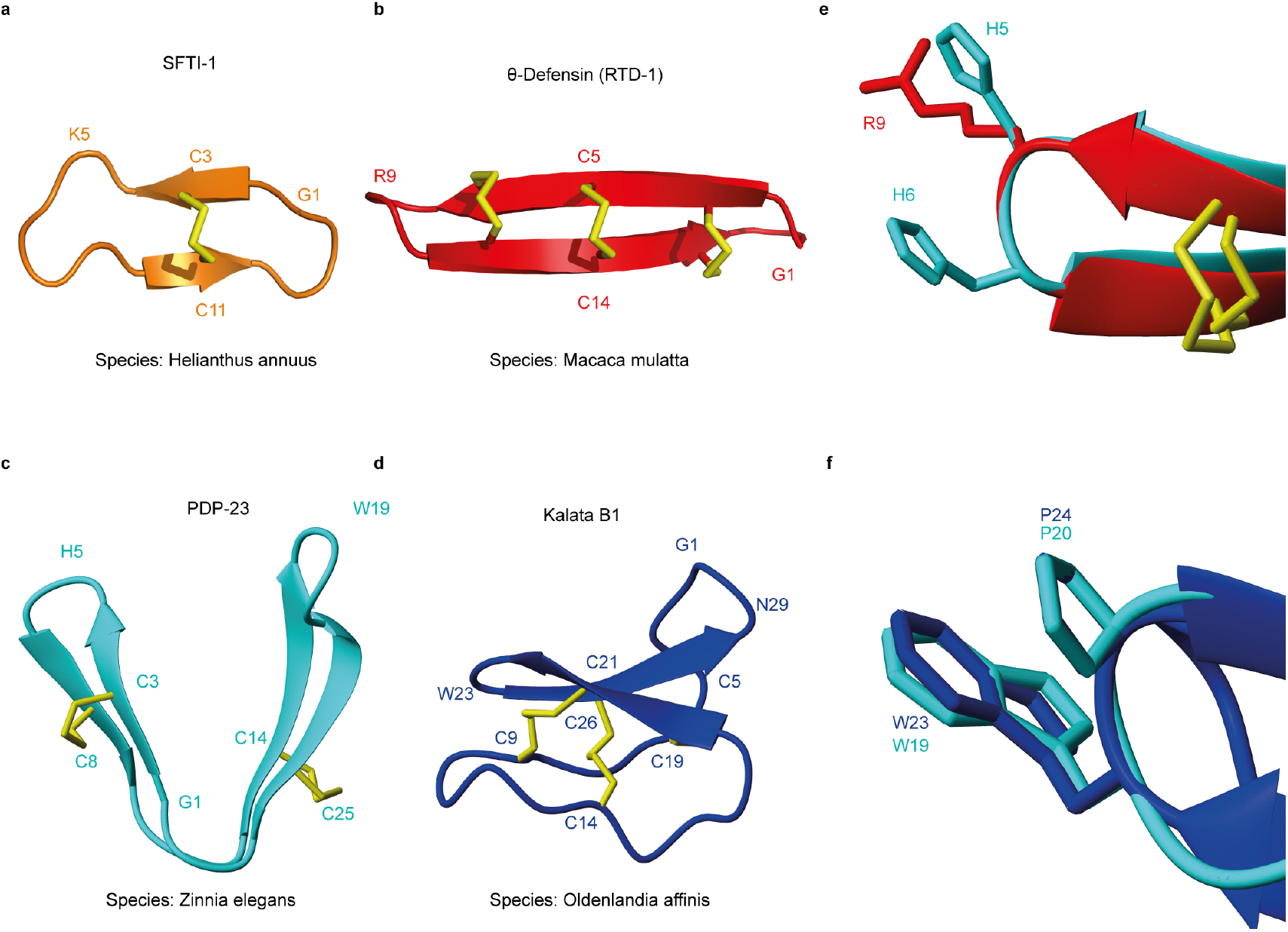
Three-dimensional structures of naturally occurring cyclic peptides. **(a)** SFTI-1 isolated from the seeds of the common sunflower *Helianthus annuus*, PDB code: 1JBL ^59^. **(b)** *θ*-defensin found in the leukocytes of rhesus macaques *Macaca mulatta*, PDB code: 2LYF ^60^. **(c)** PDP-23 isolated from the seeds of *Zinnia elegans*. **(d)** Kalata B1 isolated from the leaves of *Oldenlandia affinis*, PDB code: 1NB1 ^61^. **(e)** Overlay of type I’ turns from *θ*-defensin RTD-1 (red) and PDP-23 (cyan). Residues in the turn region are shown in stick format and are numbered according to their position in the sequence. **(f)** Overlay of type VIa1 turns from kalata B1 (blue) and PDP-23 (cyan). Residues in the turn region are shown in stick format and are numbered according to their position in the sequence.

Cyclization and the additional structural constraint imposed by disulfide bonds have been shown to provide resistance to degradation by proteases^44,45^. PDP-23 is the first naturally occurring constrained bi-disulfide peptide with a cyclic backbone, which opens up new possibilities in terms of grafting due to its different loop sizes. The smaller PDPs usually contain β-type turn loops four residues in length^16^, whereas the larger cyclotides contain six loops with the largest loop being a right turn of up to eight residues^46^. PDP-23 has a large loop between Cys^III^-Cys^IV^ containing ten amino acids that may host a larger functional loop. Hydrophobic surface patches have been described for various cyclotides including the prototypic kalata B1 and it has been shown that they are involved in membrane-binding interactions by specifically and selectively binding to PE phospholipids^48,49^. PDP-23 in contrast buries its hydrophobic side chains in the core, but is able to expose them in membrane environments. This chameleonic behavior may offer advantages in permeability, and we show that PDP-23 is able to internalize into cells without adverse toxic effects. Furthermore, the ability of PDP-23 to spatially adjust positions of turns offers new possibilities for introducing multiple functionalities in different positions.

The physiological relevance of PDP-23, and the vast majority of PDPs, in seeds remains an enigma. The potent trypsin inhibitor SFTI-1 has been proposed to have evolved as defense against gramnivore insects, given plants lack trypsin^12^. Sequences highly homologous to SFTI-1 also inhibit trypsin, but most PDPs do not show protease inhibition and PDP-23 is highly unlikely to do so based on sequence and fold. In this study PDP-23 was instead investigated as a potential scaffold for small molecule therapeutics using conjugation chemistry. Attaching 5/6-carboxy-tetramethyl-rhodamine to the PDP-23 D28K analogue yielded a conjugate that was able to demonstrate significant inhibition of P-gp by restoring the sensitivity of KB-V-1 cells to daunorubicin. 5/6-carboxy-tetramethyl-rhodamine has been previously reported to be a substrate of the P-gp transporter^30^. However, specific inhibition of P-gp has not been reported. The creation of a P-gp inhibitor out of a small molecule substrate of the transporter not only demonstrates the ability of the PDP-23 scaffold to be used as a carrier of small molecule payloads for intracellular targets, it also presents a novel way of overcoming drug resistance. This application will be particularly useful for small molecule therapeutics that are easily broken down *in vivo* as well as for chemotherapeutics where resistance to the drug is achieved by efflux. Future work will focus on modifying the PDP-23 sequence to target specific cells to direct payloads.

In conclusion we report the identification, characterization and potential applications of PDP-23, a peptide approximately twice the size of most PDPs and the first reported macrocyclic peptide with two disulfides. It folds into a remarkably optimized fold comprising extensive secondary, tertiary and quaternary structure stabilized by disulfide bonds, hydrogen bonds, and a hydrophobic core. The fold is highly resistant to proteolysis, but has a unique chameleonic character, allowing it to adapt to different environments. PDP-23 represents a novel scaffold for protein engineering, offering multiple turn sites for grafting of bioactive epitopes as well as multiple sites for the conjugation of small molecules. Importantly, unlike other commonly used scaffolds these sites are not spatially confined, PDP-23 is not toxic and PDP-23 freely internalizes into cells. We demonstrated these abilities by creating an inhibitor of P-gp and show PDP-23 is capable of potentiating chemotherapeutics against resistant cancer cells.

## Methods

### RNA-seq and transcriptome assembly for *Z. elegans*

Seeds of *Z. elegans* (Zinnia Early Wonder Mixed, Mr. Fothergill’s, Australia) were ground under liquid nitrogen with glass beads to a fine powder. Total RNA from approximately 0.3 mL of frozen tissue powder was extracted as previously described^32^. Contaminating genomic DNA was removed using DNase and the total RNA sample was further purified with a NucleoSpin RNA Clean-up kit (Macherey-Nagel). Sequencing was then performed on an Illumina HiSeq 1500 instrument as 101 bp single read runs.

The *de novo* transcriptome assemblies were performed as previously described^31,50^. We used stringent conditions for trimming and filtering of RNA-seq data. Quality trimming and filtering was done using the FASTX toolkit (hannonlab.cshl.edu/fastx_toolkit/) with raw reads trimmed with a minimum quality threshold of 30, implying a base call accuracy of 99.9%, and minimum length of 50. Trimmed reads were then filtered with a quality threshold of 30 and the percentage of bases that match the quality threshold was set to 90. *De novo* transcriptomes were assembled using CLC Genomics Workbench 8.5.1 (Qiagen). Filtered reads were assembled with five different word sizes (*i*.*e*. 23, 30, 40, 60, 64), keeping all other parameters as default. We used tBLASTn to search the transcriptome of *Z. elegans* using the *PawS1* sequence (GenBank accession FJ469150) as a query. Then the filtered raw reads were mapped back to the assembled contig encoding PDP-23 with the following default settings: Mismatch cost: 10, Insertion cost: 10, Deletion cost: 10, Length fraction: 1, Similarity fraction: 1.

### LC-MS analysis of the *Z. elegans* seed peptide extract

A small amount of dry seed peptide extract was dissolved in HPLC-grade solvent consisting of 5% (v/v) acetonitrile 0.1% formic acid (v/v) in water (Honeywell). After centrifuging for 20 min at 20,000 x *g* to remove any solids present, a 4 µL aliquot of the solution was dispensed onto a 96-well plate for LC-MS analysis. Two microliters of the sample were injected into a G4240-62010 HPLC chip (Agilent Technologies) made up of a 160 nL enrichment column (Zorbax 300SB-C18 5 µm) and a 150 mm x 75 µm analysis column (Zorbax 300SB-C18 5 µm). Separation was achieved using an 1100 series nano/capillary HPLC system (Agilent) with an elution gradient from 5% acetonitrile 0.1% formic acid to 95% acetonitrile 0.1% formic acid over 15 min at a flow rate of 0.3 µL/min. Peptides were ionized by electrospray ionization and the ions were introduced into a 6510 Q-TOF mass spectrometer (Agilent). Spectra were analyzed with MassHunter Workstation Qualitative Analysis software version B.06.00 (Agilent).

### Extraction of the macrocycle PDP-23 from *Z. elegans* seeds

*Z. elegans* seeds were purchased from either Mr. Fothergill’s (Zinnia Early Wonder Mixed) or from Royston Petrie (Lilliput Mix). To extract peptide-containing material, the seeds (100 g) were soaked in liquid nitrogen and blended to a fine meal using a food processor. A volume of 25 mL tissue powder was mixed with 160 mL of methanol containing 0.05% (v/v) trifluoroacetic acid and 160 mL of dichloromethane containing 0.05% (v/v) trifluoroacetic acid and shaken for 70 min at room temperature. After mixing, the mixture was filtered through a Whatman (No. 1) filter paper. Phase separation was achieved by adding 80 mL of 0.05% (v/v) trifluoroacetic acid in water. The top aqueous layer containing the peptide was collected, residual dichloromethane was removed under vacuum and the peptide lyophilized. Lyophilized crude mix was dissolved in 80 mL of 0.05% (v/v) trifluoroacetic acid in water, sonicated briefly and vortexed before centrifugation at 3,260 x *g* for 10 min. Crude separation was achieved by using a RP-18 Strata column (500 mg/6 mL, Phenomenex). The following buffers were used sequentially for crude separation: 10% acetonitrile 0.05% (v/v) trifluoroacetic acid (first elution); 30% acetonitrile 0.05% (v/v) trifluoroacetic acid (second elution); 90% acetonitrile 0.05% (v/v) trifluoroacetic acid (third elution). Each elution sample was analyzed by MALDI-TOF-MS to determine the monoisotopic mass of PDP-23.

### RP-HPLC purification of PDP-23

The lyophilized crude sample mix was dissolved in buffer A and split in three for further separation by RP-HPLC on a Shimadzu Prominence (Rydalmere, Australia) using an analytical Grace Vydac C18 column (250 mm x 4.6 mm, 5 µm) and an Agilent Zorbax 300SB C18 column (150 mm x 2.1 mm, 5 µm) at a 1% gradient and a flow rate of 1 mL/min and 0.3 mL/min respectively. Buffer A consisted of 0.05% TFA and buffer B consisted of 90% acetonitrile 0.05% TFA. HPLC fractions were lyophilized and stored at −20 °C.

### Sequencing of PDP-23

The PDP-23 enriched extract was dissolved in 30 µL 0.1 M ammonium bicarbonate (pH 8.3). A volume of 2 µL 200 mM dithiothreitol was added and incubated for 30 min at 60 °C to reduce the disulfide bonds. To alkylate the cysteine thiol groups, 4 µL of 500 mM iodoacetamide was added and the mixture incubated at room temperature in the dark for 10 min. To digest PDP-23 for sequencing by mass spectrometry, 1 µL of a 0.1 µg/µL stock solution of chymotrypsin (Roche Diagnostics) was added at 37 °C for 3 h. To quench the reaction prior to spectrometric analysis, 10 µL of 1% formic acid was added.

The digested PDP-23 enriched extract was analyzed by LC-MS/MS on a Shimadzu Prominence Nano HPLC (Rydalmere, Australia) coupled to a 5600 TripleTOF™ mass spectrometer (SCIEX) equipped with a nano electrospray ion source as previously described^51^.

In brief, 5 µL of digested PDP-23 enriched extract was injected and desalted. For peptide separation, a linear gradient of 2-40% solvent B over 44 min with a 500 nL/min flow rate was used, followed by a steeper gradient from 40-80% over 2 min and a hold at 80% for 2 min. The column was subsequently washed by increasing the solvent to 98% B over 0.1 min which was held at 98% for 1.9 min prior to a return to 2% B for re-equilibration (16 min). The ionspray voltage was set to 2400 V, declustering potential 100 V, curtain gas flow 25, nebuliser gas 112 and interface heater at 150 °C. The mass spectrometer was set to acquire TOF-MS data over the mass range 350-1800 for 250 ms followed by 20 full scan product ion spectra over the mass range 80-1400 with a maximum accumulation time of 250 ms in Information Dependent Acquisition mode. The 20 most intense ions observed in the TOF-MS scan exceeding a threshold of 120 counts and a charge state of +2 to +5 were set to trigger the acquisition of product ion MS/MS spectra. The data were acquired and processed using Analyst TF 1.6 software (SCIEX). ProteinPilot™ software 4.0.0.0 (SCIEX), with the paragon algorithm, was used as the MS/MS ion search engine^52^. The custom-built database contained the sequence of PDP-23 and the search parameter settings were cysteine modification with iodoacetamide and chymotrypsin as the digestion enzyme.

### Peptide synthesis

All three possible disulfide conformations for PDP-23 as well as the PDP-23 D28K analogue were assembled on 2-chlorotrityl chloride resin by Fmoc-based solid phase peptide synthesis using a CS336X peptide synthesizer (CS Bio). To facilitate selective disulfide bond formation the sidechains of Cys^II^-Cys^IV^ (conformation I), Cys^I^-Cys^IV^ (conformation II) and Cys^III^-Cys^IV^ (conformation III) were protected with acetamidomethyl groups and the other pair of Cys sidechains were protected with trityl groups. Peptides were cleaved from resin with 1% TFA in dichloromethane and lyophilized. For backbone cyclization sidechain protected peptide was dissolved in DMF to a concentration of 2 mM and one molar equivalent of *O*-(7-Azabenzotriazol-1-yl)-*N,N,N*′,*N*′-tetramethyluronium-hexafluorphosphate was added before gradually adding 10 molar equivalents of *N,N*-diisopropylethylamine. The solution was stirred for 6 h at room temperature before being lyophilized. Final peptide cleavage was achieved by dissolving the lyophilized peptide in 10 mL of TFA, triisopropylsilane, 2,2′-(ethylenedioxy)diethanethiol and water in a ratio of 97:1:1:1, respectively. After stirring for 2 h at room temperature, TFA was removed under vacuum and the peptide was precipitated in ice-cold ether, filtered, and dissolved in 50% acetonitrile with 0.05% TFA. Residual ether was removed under vacuum and the peptide lyophilized. The crude peptide was purified by RP-HPLC on a Shimadzu Prominence (Rydalmere, Australia) using a preparative Phenomenex Jupiter C18 column (250 mm x 21.2 mm, 10 µm) at a 1% gradient and a flow rate of 8 mL/min. Electrospray ionization mass spectrometry confirmed the molecular mass of the reduced peptides. Peptides were lyophilized before oxidization.

To form the first disulfide bond between unprotected cysteines, peptide (0.25 mg/mL) was dissolved in 0.1 M ammonium bicarbonate buffer (pH 8.3) with 2 mM reduced glutathione, stirred for 24 h at room temperature and then purified by RP-HPLC on a Shimadzu Prominence (Rydalmere, Australia) using a semi-preparative Grace Vydac C18 column (250 mm x 10 mm, 10 µm) at a 1% gradient and a flow rate of 3 mL/min. MALDI-TOF-mass spectrometry confirmed the molecular mass of the partly oxidized peptides. Peptides were lyophilized.

The second disulfide bond was formed between the acetamidomethyl-protected cysteine residues by dissolving the peptide in water containing 0.05% TFA at a concentration of 0.5 mg/mL. A saturated iodine solution was slowly added until the mixture became light yellow in color and this was incubated at 23 °C overnight in the dark. The reaction was quenched by adding ascorbic acid until the mixture became colorless again. The fully oxidized peptides were purified by RP-HPLC on a Shimadzu Prominence (Rydalmere, Australia) using a semi-preparative Grace Vydac C18 column (250 mm x 10 mm, 10 µm) at a 1% and 0.5% gradient and a flow rate of 3 mL/min. MALDI-TOF-mass spectrometry confirmed the molecular mass of the fully oxidized peptides to have a theoretical molecular mass of [M+H]^+^ 3109.24.

### Peptide labelling

The PDP-23 D28K analogue was labelled with ATTO 488 (Sigma-Aldrich, Missouri, USA) as follows: PDP-23 D28K (5 mg) was dissolved in 0.1 M sodium bicarbonate (pH 8.3) to a concentration of 2 mg/mL. Then a pre-prepared solution of ATTO 488 NHS ester (10 mg/ml in DMSO) was added to the peptide solution to obtain a ratio of 2:1 (dye:peptide). The solution was stirred at 23 °C for 2 h in the dark. The conjugation solution was diluted 10-fold and RP-HPLC was performed to filter and purify the labelled peptide from salt, excess dye, and side-reaction products. The PDP-23-ATTO488 conjugate was successfully collected showing the correct *m/z* of 924.89 [M + 4H]^4+^ and 1232.86 [M + 3H]^3+^ indicating the correct total mass of 3695.58 Da. The conjugation of 5/6-carboxy-tetramethyl-rhodamine to the PDP-23 D28K analogue was conducted as follows. PDP-23 D28K (∼5 mg) was dissolved in 0.1 M sodium phosphate to a concentration of 1 mg/mL and pH adjusted to 8.75. To this peptide solution a pre-prepared solution containing 10 molar equivalents of NHS-5/6-carboxy-tetramethyl-rhodamine at a concentration of 10 mg/mL in DMSO was added. This solution was stirred at room temperature overnight, protected from light. The conjugation solution was diluted 10-fold and RP-HPLC was performed to desalt, remove excess dye and purify the peptide-drug conjugate from side-reaction products. The PDP-23-5/6-carboxy-tetramethyl-rhodamine was successfully collected showing the correct *m/z* of 884.8 [M + 4H]^4+^ and 1179.6 [M + 3H]^3+^ indicating the correct total mass of 3536.49 Da.

### NMR spectroscopy

For analysis in water PDP-23 was dissolved in 550 µL 90:10 H_2_O/D_2_O (v/v) at pH ∼4 to a final concentration of 1.25 mg/mL. Acetonitrile samples were prepared with ∼ 1 mg/mL PDP-23 in 550 µL of 80:20 (v/v) H_2_O/CD_3_CN. For studies in micelles PDP-23 (∼1 mg) was dissolved in 550 µL of 100 mM deuterated SDS or 12 mM deuterated DPC in 90:10 H_2_O/D_2_O. Standard homonuclear 2D datasets were recorded at 298 K on 600, 700 or 900 MHz Bruker Avance III spectrometers equipped with cryoprobes, and processed using Topsin 4.0.3 (Bruker Biospin). Mixing times of 80 ms for the TOCSY and 200 ms for the NOESY were used. Additional ^13^C and ^15^N HSQC (Heteronuclear Single Quantum Coherence Spectroscopy) spectra were recorded at natural abundance. Temperature coefficients were determined at temperatures ranging from 288 to 308 K. Temperature stability studies were performed at temperatures ranging from 298 to 363 K. ^13^C HSQC data were recorded in 80:20 D_2_O/CD_3_CN (v/v) or 100% D_2_O to minimize overlap of Hα-Cα resonances with the residual water. Data were referenced to DSS at 0.0 ppm.

### Spectral assignment and structure calculations

The recorded TOCSY, NOESY, ^13^C HSQC and ^15^N HSQC spectra were used for manual chemical shift assignments using sequential assignment strategies in CARA. The structure of PDP-23 was calculated from inter-proton distance restraints generated from the peak volumes of NOESY cross peaks and dihedral ϕ (C_-1_-N-CA-C) and ψ (N-CA-C-N_+1_) backbone angles were generated by Torsion Angle Likelihood Obtained from Shift and sequence similarity (TALOS-N). A set of 50 initial structures was generated using CYANA 3.98^53^ and final structures were annealed and refined in explicit water within CNS 1.21 using protocols from the RECOORDscript database^54,55^. From the 50 calculated structures, a set of 20 structures with no violations, low energy and best MolProbity score was chosen^56^. In the case of the symmetrical homodimer structure of PDP-23, the symmetrical homodimer function in CYANA 3.98 was used to generate 50 dimeric structures wherein each monomer was restrained to the other based on the determined intermolecular NOEs observed in the NOESY spectra. These structures were then further refined in explicit water using CNS 1.21.

### *In vitro* serum stability

PDP-23 was mixed with human pooled male serum (Sigma-Aldrich) to a final concentration of 50 µg/mL and incubated at 37 °C. Aliquots (100 µL) were taken at different time points (0, 2, 4, 6 and 24 h) and quenched with 100 mM ammonium acetate, pH 3 (900 µL), followed by incubation on ice for 30 min. To separate the peptide from the serum components, Oasis HLB 3 cc 60 mg cartridges (Waters) were activated with 6 mL methanol washes, preconditioned with 3 mL 70% acetonitrile, 1% formic acid and equilibrated with 3 mL 1% formic acid. The serum sample was then loaded onto the column and further washed with 3 mL 5% acetonitrile, 1% formic acid before elution with 35% acetonitrile, 1% formic acid^57^. Samples were lyophilized and re-dissolved in 100 µL 1% formic acid before LC-MS/MS analysis (AB Sciex-API2000) and quantification (AB Sciex-MultiQuant). The serum stability tests were performed in triplicate and peptide stability was calculated using a non-linear fit of one phase decay in GraphPad Prism 6.

### Simulated intestinal fluid stability

Simulated intestinal fluid (SIF) was prepared according to U.S Pharmacopeia specification. Briefly, 68 mg KH_2_PO_4_ was dissolved in 250 µL MilliQ water. To this solution, 770 µL 0.2 M NaOH and 5 ml of MiliQ water were added, mixed with 100 µg of pancreatin from porcine pancreas 8x U.S.P (Sigma Aldrich). This mixture was adjusted to pH 6.8 ± 0.1 with 0.2 M HCl or 0.2 M NaOH and diluted with water to a final volume of 10 mL. SIF was pre-incubated for 15 min at 37 °C, before addition of 50 µg peptide to 500 µL SIF. Samples were taken at different time points (0, 0.083, 0.33, 1, 2, 4, 6 and 24 h). Each SIF aliquot (50 µL) was quenched with 50 µL 4% TFA. Samples were analyzed by RP-HPLC on an analytical Grace Vydac C18 column (2.1 mm x 150 mm, 5 µm) using a linear aqueous acetonitrile gradient containing 0.05% TFA at a flow rate of 0.3 mL/min. PDP-23 was detected by recording the sample absorbance at 214 nm and quantified by the peak area relative to the peak area at time point 0 min.

### Simulated gastric fluid stability

Simulated gastric fluid (SGF) was prepared according to U.S Pharmacopeia specification. In brief, 20 mg of NaCl and 32 mg of pepsin from porcine gastric mucosa (Sigma Aldrich) were dissolved in 10 mL of 70 µL of HCl in water (final pH 1.2). SGF was pre-incubated for 15 min at 37 °C, before addition of 50 µg peptide to 500 µL SGF. Samples were taken at different time points (0, 0.083, 0.33, 0.66, 1, 4 and 24 h). Each SGF aliquot (50 µL) was quenched with 50 µL 0.2 M Na_2_CO_3_. Samples were analyzed by RP-HPLC on an analytical Grace Vydac C18 column (2.1 mm x150 mm, 5 µm) using a linear aqueous acetonitrile gradient containing 0.05% TFA and a flow rate of 0.3 mL/min. PDP-23 was detected by recording the sample absorbance at 214 nm and quantified by the peak area relative to the peak area at time point 0 min.

### Cell Culture for PDP-23 toxicity assays

Human cervical cancer (HeLa) cells were used to assess the toxicity of PDP-23. The medium used for culture was Eagle’s Minimum Essential Medium supplemented with 10% fetal calf serum (FCS), 50 U/mL penicillin and 50 μg/mL streptomycin. Cells were incubated at 37 °C in 5% CO_2_. PDP-23 was evaluated for cellular toxicity at concentrations up to 32 µM using the standard 3-(4,5,-dimethyl-2-thazolyl)-2,5-diphenyl-2H-tetrazolium bromide (MTT) assay^58^.

### Cell uptake assays

The uptake of PDP-23 was visualized using confocal microscopy. HeLa cells were seeded on sterile coverslips, placed into a 24-well culture plate at a density of 10^4^ cells/well and incubated overnight. The medium was then removed and the cells incubated in fresh medium containing ATTO 488-labeled PDP-23-D28K at a concentration of 2 µM for 2 h. This medium was then removed and the cells washed with cold PBS. The cells were then treated with fresh medium containing 5 µg/mL wheat germ agglutinin conjugated with Alexa-633 and kept at 4 °C for 30 min. The cells were then washed with cold PBS prior to fixing the cells by incubating them with 4% paraformaldehyde for 15 min at 4 °C. The cells were again washed with cold PBS and the sample mounted in an antifade medium (VECTASHIELD® Antifade Mounting Medium). Microscopy was then performed using a Leica DMi8 SP8 inverted microscope.

### Cell culture and chemical preparation for PDP-23/rhodamine conjugate toxicity assays

KB-3-1 (sensitive) and KB-V-1 (resistant) cells were grown on poly-l-lysine-coated T75 flasks at 37 °C with 5% CO_2_ and maintained in RPMI 1640 medium, supplemented with 10% FCS, 2 mM glutamine and 100 U/mL & 100 µg/mL penicillin/streptomycin. Daunorubicin and verapamil were dissolved in DMSO and diluted in RPMI medium to produce working stocks, with the final solvent concentration not exceeding 1%. PDP-23-5/6-carboxy-tetramethyl-rhodamine (159 µg) was reconstituted in 1.5 mL of RPMI to a concentration of 30 µM, which was further diluted to produce working stocks.

### Lysate preparation and western blot

KB-3-1-and KB-V-1 cells were plated at a density of 300,000 cells per well of a 12-well plate in 1 mL of the RPMI medium and incubated overnight. Cells were then lysed in Laemmli’s buffer and heated for 5 min at 95 °C. Lysates were electrophoresed at 200 V for 45 min on a 7.5% SDS-acrylamide gel and transferred at 350 mA for 1 h to a nitrocellulose membrane. Blocking was performed for 1 h at RT in 5% skim milk/PBS with 0.05% Tween-20 (PBST). The membrane was washed thrice with PBST for 15 min and rocked overnight at 4 °C with P-gp primary antibody. The membrane was washed again and rocked with rabbit horseradish peroxidase-conjugated secondary antibody for 1 h at RT. After another set of washes, P-gp expression was detected via ECL. The loading control was generated by repeating the steps post blocking using an α-tubulin primary antibody and a mouse horseradish peroxidase-conjugated secondary antibody.

### Daunorubicin sensitivity determination

Cells were seeded in a 96-well plate at a density of 10,000 cells per well in 100 µL of RPMI medium and incubated overnight at 37 °C. Cells were treated with DMSO or one of nine concentrations of daunorubicin for 24 h. Medium was removed and cells incubated with the DNA-binding fluorescent dye reagent (CyQuant NF Proliferation assay kit) in Hank’s Balanced Salt Solution, supplemented with 20 mM 4-(2-hydroxyethyl)-1-piperazineethanesulfonic acid and 35 mg/mL NaHCO_3_, for 30 min at 37 °C. Fluorescence was measured at 485_ex_/520_em_. Data were normalized to the untreated control. For PDP-23-5/6-carboxy-tetramethyl-rhodamine conjugate toxicity assays, cells were plated in a 96-well plate as described above. Cells were treated with positive controls of 30 µM verapamil supplemented with 1 µM daunorubicin, as well as daunorubicin 1 µM alone. Cells were additionally treated with PDP-23-5/6-carboxy-tetramethyl-rhodamine at concentrations of 1, 3, 10 or 30 µM supplemented with 1 µM daunorubicin. Negative controls included untreated cells as well as cells treated with 30 µM verapamil only. Toxicity was assessed using the CyQuant NF Proliferation assay protocol as above. Data were normalized to the untreated control.

## Acknowledgments

This work, B.F. and G.V. were supported by Australian Research Council (ARC) grants DP120103369 and DP190102058 to J.S.M and K.J.R. C.D.P. was supported by a UQ Postgraduate Research Award. M.F.F. was supported by an Australian Postgraduate Award and a Bruce and Betty Green Postgraduate Research Scholarship. C.E.M. was supported by an Australian Postgraduate Award. J.Z., A.S.J. and F.H. were supported by International Postgraduate Research Scholarships and an Australian Postgraduate Award. K.J.R. and J.S.M. were supported by ARC Future Fellowships FT130100890 and FT120100013 respectively.

## Author Contributions

J.S.M. and K.J.R. conceived the study; J.S.M. extracted RNA, A.S.J. and J.Z. performed transcriptomics, M.F.F. performed LC-MS to confirm presence of PDP-23 in seed extract. C.D.P., B.F., A.S., F.H. and R.J.C. performed peptide synthesis. C.D.P., B.F., C.E. and K.J.R. analysed NMR data and calculated structures. A.S. performed stability assays. F.H. performed cell toxicity and uptake assays. C.D.P. performed drug conjugation to scaffold. C.E.M. and R.F.M. performed P-gp assays. All authors analyzed data; C.D.P., B.F., J.S.M. and K.J.R. wrote the manuscript with contributions from all authors.

## References

1 Bhardwaj, G. et al. Accurate de novo design of hyperstable constrained peptides. Nature 538, 329–335, (2016).

2 Correnti, C. E. et al. Screening, large-scale production and structure-based classification of cystine-dense peptides. Nat. Struct. Mol. Biol. 25, 270–278, (2018).

3 Wang, C. K. & Craik, D. J. Designing macrocyclic disulfide-rich peptides for biotechnological applications. Nat. Chem. Biol. 14, 417–427, (2018).

4 González, C. et al. Bacteriocin AS-48, a microbial cyclic polypeptide structurally and functionally related to mammalian NK-lysin. PNAS 97, 11221–11226, (2000).

5 Vetter, J. Toxins of Amanita phalloides. Toxicon. 36, 13–24, (1998).

6 Tang, Y.-Q. et al. A cyclic antimicrobial peptide produced in primate leukocytes by the ligation of two truncated α-defensins. Science 286, 498–502, (1999).

7 Saether, O. et al. Elucidation of the primary and three-dimensional structure of the uterotonic polypeptide kalata B1. Biochemistry 34, 4147–4158, (1995).

8 Franke, B., Mylne, J. S. & Rosengren, K. J. Buried treasure: biosynthesis, structures and applications of cyclic peptides hidden in seed storage albumins. Nat. Prod. Rep. 35, 137–146, (2018).

9 Luckett, S. et al. High-resolution structure of a potent, cyclic proteinase inhibitor from sunflower seeds J. Mol. Biol. 290, 525–533, (1999).

10 Mylne, J. S. et al. Albumins and their processing machinery are hijacked for cyclic peptides in sunflower. Nat. Chem. Biol. 7, 257–259, (2011).

11 Bernath-Levin, K. et al. Peptide macrocyclization by a bifunctional endoprotease. Chem. Biol. 22, 571–582, (2015).

12 Elliott, A. G. et al. Evolutionary origins of a bioactive peptide buried within preproalbumin. Plant Cell 26, 981–995, (2014).

13 Jayasena, A. S. et al. Stepwise evolution of a buried inhibitor peptide over 45 My. Mol. Biol. Evol. 34, 1505–1516, (2017).

14 Fisher, M. F. et al. A family of small, cyclic peptides buried in preproalbumin since the Eocene epoch. Plant Direct 2, e00042, (2018).

15 Franke, B. et al. Diverse cyclic seed peptides in the Mexican zinnia (Zinnia haageana). Biopolymers 106, 806–817, (2016).

16 Elliott, A. G. et al. Natural structural diversity within a conserved cyclic peptide scaffold. Amino Acids 49, 103–116, (2017).

17 Clark, R. J. et al. Engineering stable peptide toxins by means of backbone cyclization: stabilization of the alpha-conotoxin MII. PNAS 102, 13767–13772, (2005).

18 Clark, R. J. et al. The engineering of an orally active conotoxin for the treatment of neuropathic pain. Angew. Chem. 49, 6545–6548, (2010).

19 Gaurav, B. et al. Accurate de novo design of hyperstable constrained peptides. Nature 538, 329–335, (2016).

20 Ji, Y. et al. In vivo activation of the p53 tumor suppressor pathway by an engineered cyclotide. J. Am. Chem. Soc. 135, 11623–11633, (2013).

21 D’Souza, C. et al. Using the MCoTI-II cyclotide scaffold to design a stable cyclic peptide antagonist of SET, a protein overexpressed in human cancer. Biochemistry 55, 396–405, (2016).

22 Wang, C. K. et al. Molecular grafting onto a stable framework yields novel cyclic peptides for the treatment of multiple sclerosis. ACS chem. biol. 9, 156–163, (2014).

23 Chan, L. Y. et al. Engineering pro-angiogenic peptides using stable, disulfide-rich cyclic scaffolds. Blood 118, 6709–6717, (2011).

24 de Veer, S. J. et al. Engineered protease inhibitors based on sunflower trypsin inhibitor-1 (SFTI-1) provide insights into the role of sequence and conformation in Laskowski mechanism inhibition. Biochem. J. 469, 243–253, (2015).

25 Li, C. Y. et al. Amino acid scanning at P5’ within the Bowman-Birk inhibitory loop reveals specificity trends for diverse serine proteases. J. Med. Chem. 62, 3696–3706, (2019).

26 He, R., Finan, B., Mayer, J. P. & DiMarchi, R. D. Peptide conjugates with small molecules designed to enhance efficacy and safety. Molecules 24, 1855, (2019).

27 de Jong, M. et al. Somatostatin receptor-targeted radionuclide therapy of tumors: Preclinical and clinical findings. Semin. Nucl. Med. 32, 133–140, (2002).

28 Westphalen, S. et al. Receptor mediated antiproliferative effects of the cytotoxic LHRH agonist AN-152 in human ovarian and endometrial cancer cell lines. Int. J. Oncol. 17, 1063–1069, (2000).

29 Régina, A. et al. Antitumour activity of ANG1005, a conjugate between paclitaxel and the new brain delivery vector Angiopep-2. Br. J. Pharmacol. 155, 185–197, (2008).

30 Loetchutinat, C., Saengkhae, C., Marbeuf-Gueye, C. & Garnier-Suillerot, A. New insights into the P-glycoprotein-mediated effluxes of rhodamines. Eur. J. Biochem. 270, 476–485, (2003).

31 Jayasena, A. S. et al. Next generation sequencing and de novo transcriptomics to study gene evolution. Plant Methods 10, 34, (2014).

32 Mylne, J. S. et al. Cyclic peptides arising by evolutionary parallelism via asparaginyl-endopeptidase-mediated biosynthesis. Plant Cell 24, 2765–2778, (2012).

33 Schroeder, C. I. & Rosengren, K. J. in Snake and Spider Toxins: Methods and Protocols (ed Avi Priel) 129–162 (Springer US, 2020).

34 Duplâtre, G., Ferreira Marques, M. F. & da Graça Miguel, M. Size of sodium dodecyl sulfate micelles in aqueous solutions as studied by positron annihilation lifetime spectroscopy. J. Phys. Chem. 100, 16608–16612, (1996).

35 Sikorska, E. et al. Thermodynamics, size, and dynamics of zwitterionic dodecylphosphocholine and anionic sodium dodecyl sulfate mixed micelles. J. Therm. Anal. Calorim. 123, 511–523, (2016).

36 Gottesman, M. M., Fojo, T. & Bates, S. E. Multidrug resistance in cancer: role of ATP– dependent transporters. Nat. Rev. Cancer. 2, 48–58, (2002).

37 Nanayakkara, A. K. et al. Targeted inhibitors of P-glycoprotein increase chemotherapeutic-induced mortality of multidrug resistant tumor cells. Sci. Rep. 8, (2018).

38 Göransson, U. & Craik, D. J. Disulfide mapping of the cyclotide kalata B1. Chemical proof of the cyclic cystine knot motif. J. Biol. Chem. 278, 48188–48196, (2003).

39 Craik, D. J., Daly, N. L., Bond, T. & Waine, C. Plant cyclotides: A unique family of cyclic and knotted proteins that defines the cyclic cystine knot structural motif. J. Mol. Biol. 294, 1327–1336, (1999).

40 Pallaghy, P. K., Nielsen, K. J., Craik, D. J. & Norton, R. S. A common structural motif incorporating a cystine knot and a triple-stranded β-sheet in toxic and inhibitory polypeptides. Protein Sci. 3, 1833–1839, (1994).

41 Norton, R. S. & Pallaghy, P. K. The cystine knot structure of ion channel toxins and related polypeptides. Toxicon. 36, 1573–1583, (1998).

42 Daly, N. L. et al. Retrocyclin-2: structural analysis of a potent anti-HIV theta-defensin. Biochemistry 46, 9920–9928, (2007).

43 Rosengren, K. J., Daly, N. L., Harvey, P. J. & Craik, D. J. The self-association of the cyclotide kalata B2 in solution is guided by hydrophobic interactions. Biopolymers 100, 453–460, (2013).

44 Colgrave, M. L. & Craik, D. J. Thermal, chemical, and enzymatic stability of the cyclotide kalata B1: the importance of the cyclic cystine knot. Biochemistry 43, 5965–5975, (2004).

45 Wang, J., Yadav, V., Smart, A. L., Tajiri, S. & Basit, A. W. Toward oral delivery of biopharmaceuticals: an assessment of the gastrointestinal stability of 17 peptide drugs. Mol. Pharm. 12, 966–973, (2015).

46 Hernandez, J. F. et al. Squash trypsin inhibitors from Momordica cochinchinensis exhibit an atypical macrocyclic structure. Biochemistry 39, 5722–5730, (2000).

47 Barry, D. G., Daly, N. L., Bokesch, H. R., Gustafson, K. R. & Craik, D. J. Solution structure of the cyclotide palicourein: implications for the development of a pharmaceutical framework. Structure 12, 85–94, (2004).

48 Henriques, S. T. et al. Decoding the membrane activity of the cyclotide kalata B1: the importance of phosphatidylethanolamine phospholipids and lipid organization on hemolytic and anti-HIV activities. J. Biol. Chem. 286, 24231–24241, (2011).

49 Henriques, S. T. et al. Phosphatidylethanolamine binding is a conserved feature of cyclotide-membrane interactions. J. Biol. Chem. 287, 33629–33643, (2012).

50 Garcia-Seco, D., Zhang, Y., Gutierrez-Mañero, F. J., Martin, C. & Ramos-Solano, B. RNA-Seq analysis and transcriptome assembly for blackberry (Rubus sp. Var. Lochness) fruit. BMC Genom. 16, (2015).

51 Franke, B., Colgrave, M. L., Mylne, J. S. & Rosengren, K. J. Mature forms of the major seed storage albumins in sunflower: A mass spectrometric approach. J. Proteomics 147, 177–186, (2016).

52 Shilov, I. V. et al. The paragon algorithm, a next generation search engine that uses sequence temperature values and feature probabilities to identify peptides from tandem mass spectra. Mol. Cell Proteomics 6, 1638–1655, (2007).

53 Güntert, P. Automated NMR structure calculation with CYANA. Methods Mol. Biol. 278, 353–378, (2004).

54 Brunger, A. T. Version 1.2 of the Crystallography and NMR system. Nat. Protoc. 2, 2728–2733, (2007).

55 Nederveen, A. J. et al. RECOORD: a recalculated coordinate database of 500+ proteins from the PDB using restraints from the BioMagResBank. Proteins 59, 662–672, (2005).

56 Chen, V. B. et al. MolProbity: all-atom structure validation for macromolecular crystallography. Acta Crystallogra. D 66, 12–21, (2010).

57 Hossain, M. A., Haugaard-Kedström, L. M., Rosengren, K. J., Bathgate, R. A. & Wade, J. D. Chemically synthesized dicarba H2 relaxin analogues retain strong RXFP1 receptor activity but show an unexpected loss of in vitro serum stability. Org. Biomol. Chem. 13, 10895–10903, (2015).

58 van Meerloo, J., Kaspers, G. J. & Cloos, J. Cell sensitivity assays: the MTT assay. Methods. Mol. Biol. 731, 237–245, (2011).

59 Korsinczky, M. L. et al. Solution structures by ^1^H NMR of the novel cyclic trypsin inhibitor SFTI-1 from sunflower seeds and an acyclic permutant. J. Mol. Biol. 311, 579–579, (2001).

60 Conibear, A. C., Rosengren, K. J., Harvey, P. J. & Craik, D. J. Structural characterization of the cyclic cystine ladder motif of θ-defensins. Biochemistry 51, 9718–9726, (2012).

61 Rosengren, K. J., Daly, N. L., Plan, M. R., Waine, C. & Craik, D. J. Twists, knots, and rings in proteins. Structural definition of the cyclotide framework. J. Biol. Chem. 278, 8606–8616, (2003).

