## Supplementary Information for "A Chameleonic Macrocyclic Peptide with Drug Delivery Applications"

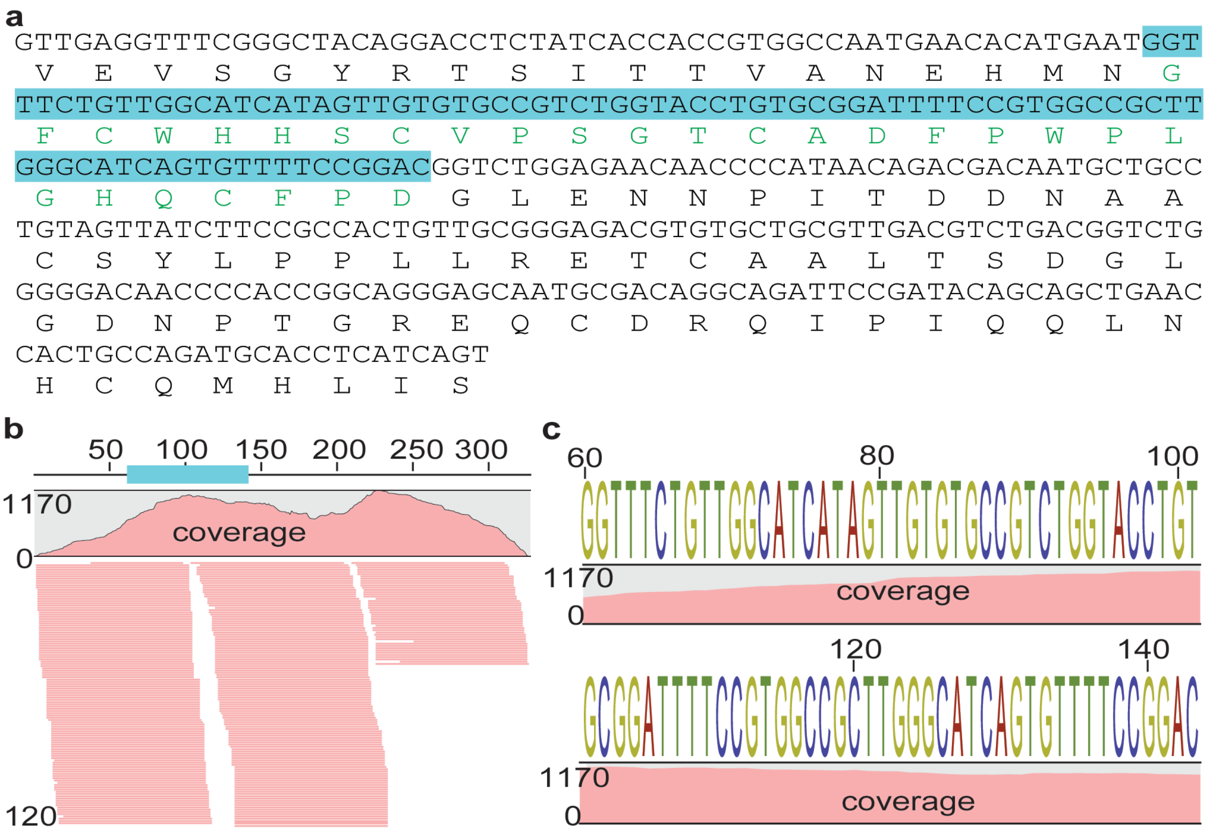


**Supplementary Figure 1: Contig sequence that encodes PDP-23 and sequence support.**

**(a)** Sequence assembled *de novo* from RNA-seq of *Zinnia elegans* seeds that encodes PDP-23 (cyan highlight) and its translated sequence. **(b)** Mapping RNA-seq reads to the contig shows strong support for the sequence with the average depth of coverage being 696 reads. **(c)** The sequence logo generated from mapped reads shows high confidence for the PDP-23 sequence with no evidence for sequence polymorphism.

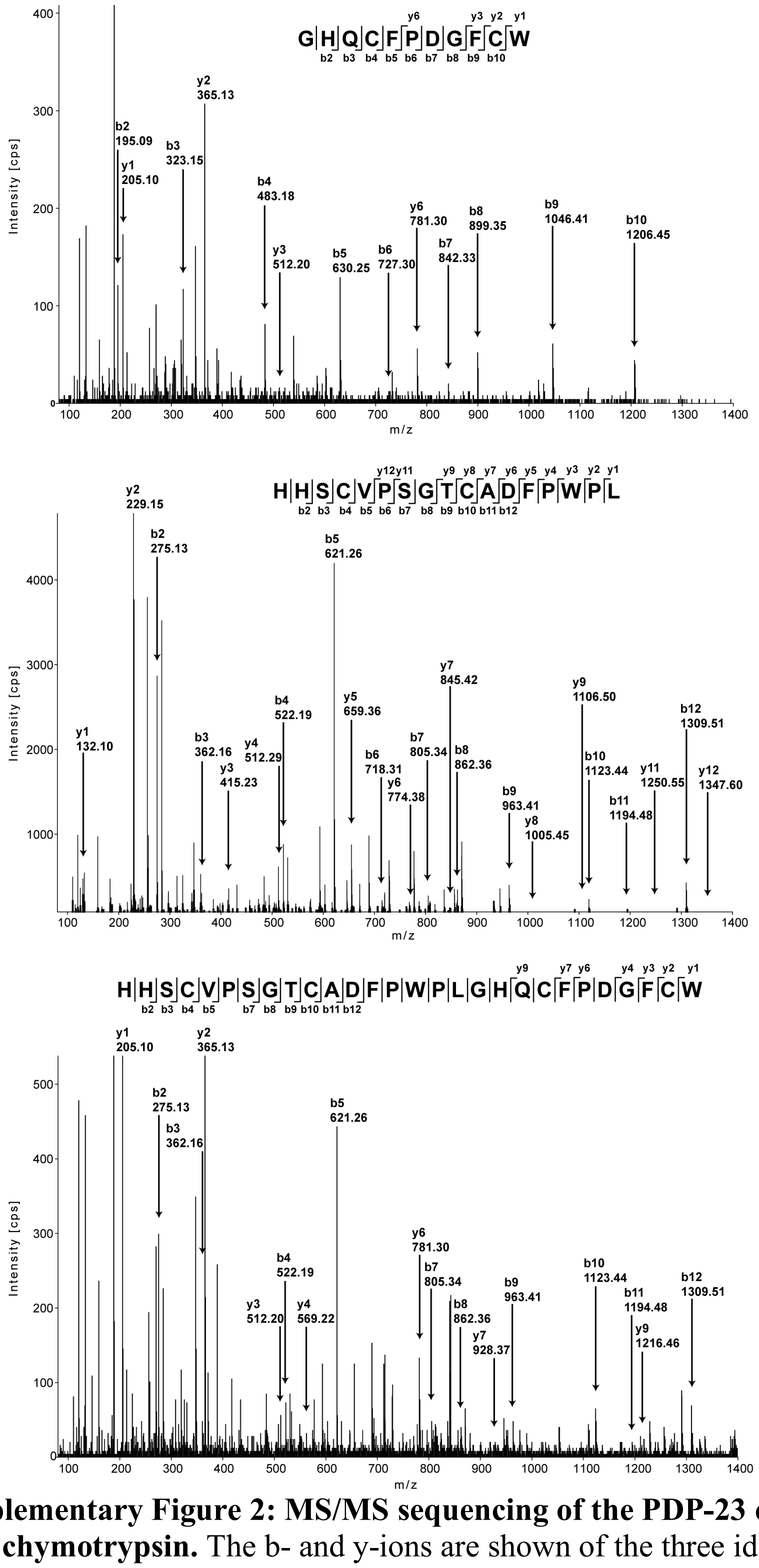


**Supplementary Figure 2: MS/MS sequencing of the PDP-23 enriched extract digested with chymotrypsin.** The b- and y-ions are shown of the three identified fragments after the PDP-23 enriched extract was reduced, alkylated and digested with chymotrypsin.

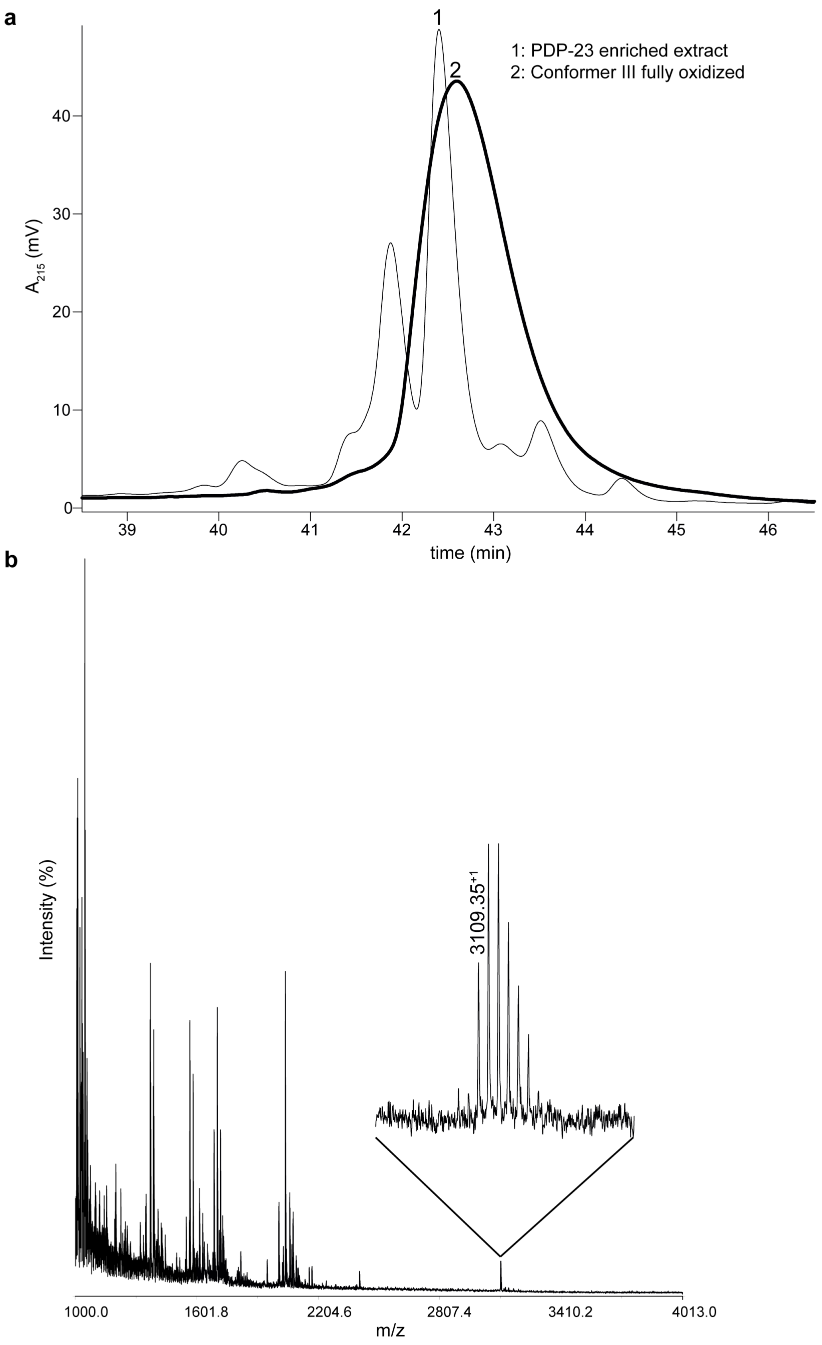


**Supplementary Figure 3: Analytical HPLC profile of conformer III and PDP-23 enriched extract. (a)** Analytical HPLC trace of conformer III highlighted in a solid bold black line overlaid with a trace of PDP-23 enriched extract. **(b)** MALDI-TOF mass spectrum showing the mass-to-charge ratio (*m/z*) of PDP-23 alongside many other masses in the PDP-23 enriched extract.

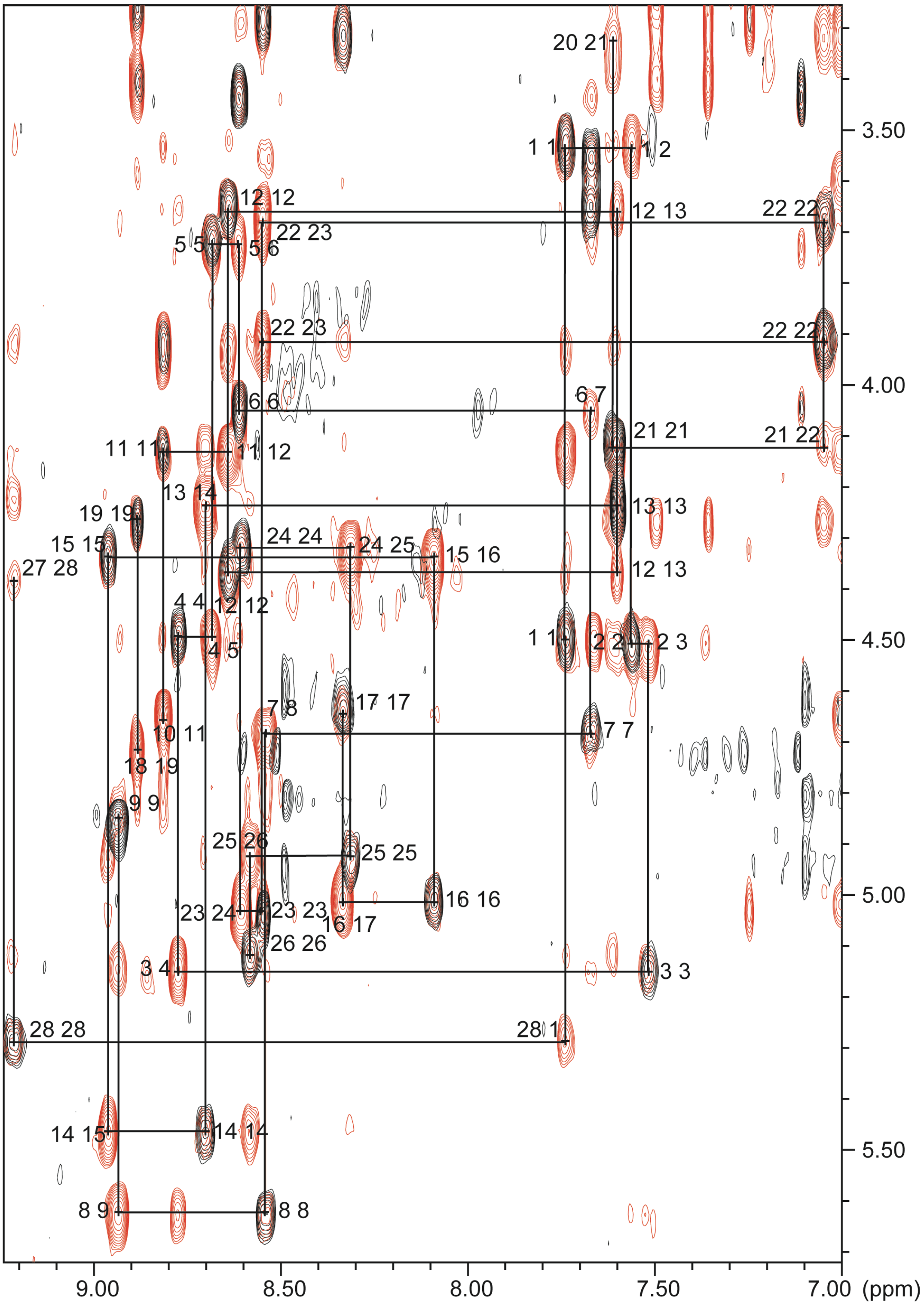


**Supplementary Figure 4: Sequential assignment of the NMR data.** The 'sequential walk' in the 'fingerprint' Hα-HN region of the NMR spectra is shown for PDP-23. Data were recorded in 90:10 ($H_2O/D_2O$) at 600 MHz and 298 K. The TOCSY spectrum (80 ms mixing time; black), which contain only intra-residual connections is superimposed on the NOESY spectrum (200 ms mixing time; red), which contain both intra- and inter-residual connections. The sequential walk is only broken by proline residues at positions 10, 18, 20, 27.

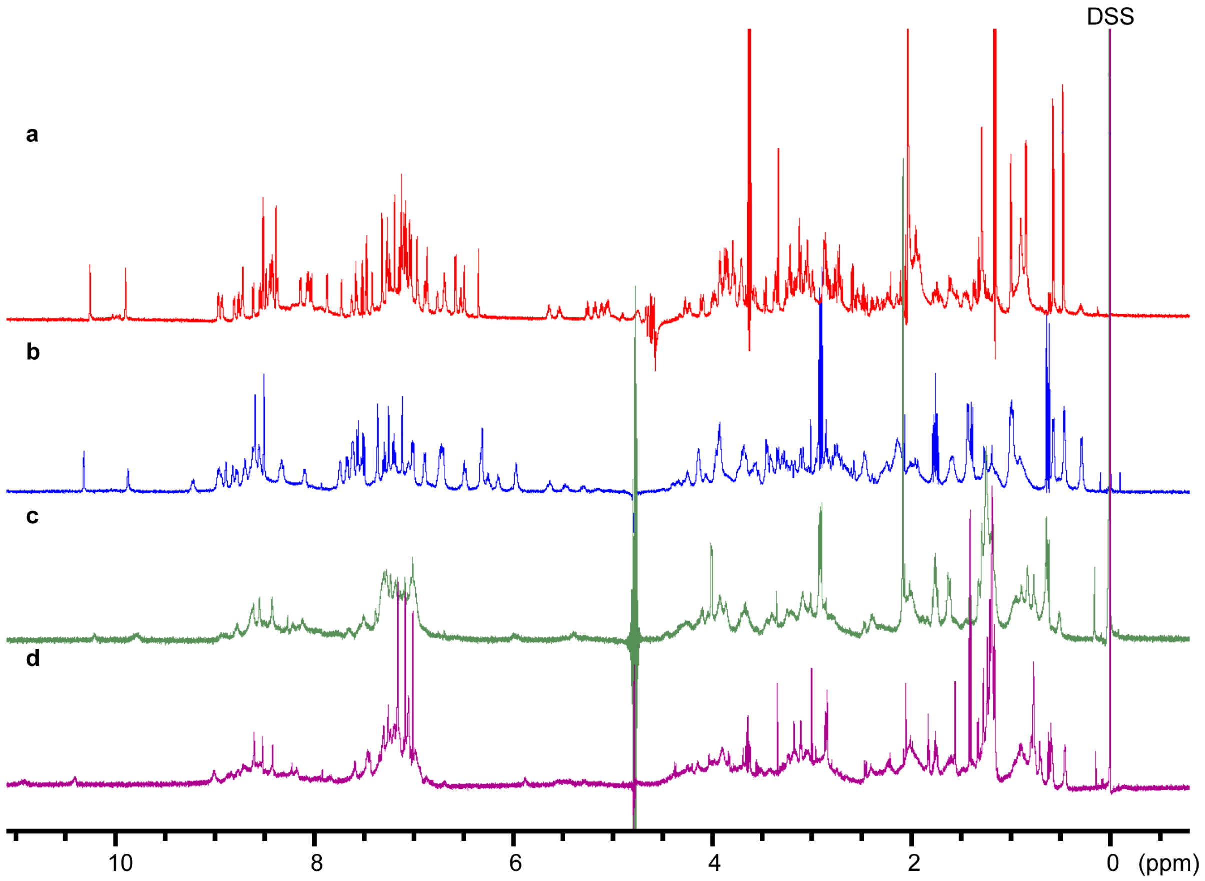


**Supplementary Figure 5: One-dimensional $^{1}$H NMR spectra of PDP-23 in different environments. (a)** PDP-23 in 80:20 ($H_2O/CD_3CN$), **(b)** PDP-23 in 90:10 ($H_2O/D_2O$), **(c)** PDP-23 when exposed to SDS micelles, **(d)** PDP-23 when exposed to DPC micelles. All spectra were recorded at 298K, and are aligned to DSS at 0 ppm.

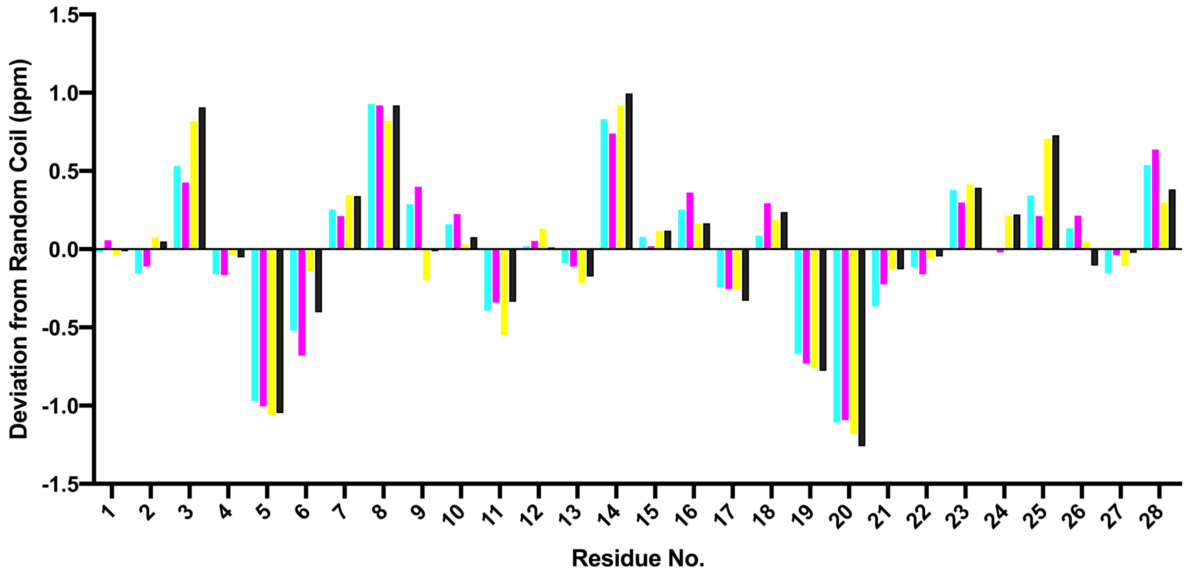


**Supplementary Figure 6: Secondary Hα chemical shifts of PDP-23 in different environments.** Secondary shifts, the difference between observed chemical shifts and chemical shifts observed in short unstructured random coil peptides, are sensitive indicators of secondary structure. Negative values are typical of helical or turn structure, while positive values suggest extended sheet structure. Columns are color coded according to conditions; 80:20 $H_2O/CD_3CN$ (cyan), 90:10 ($H_2O/D_2O$) (magenta), SDS micelles (yellow), DPC micelles (black). The secondary shifts are remarkably similar, confirming the secondary structure is largely the same.

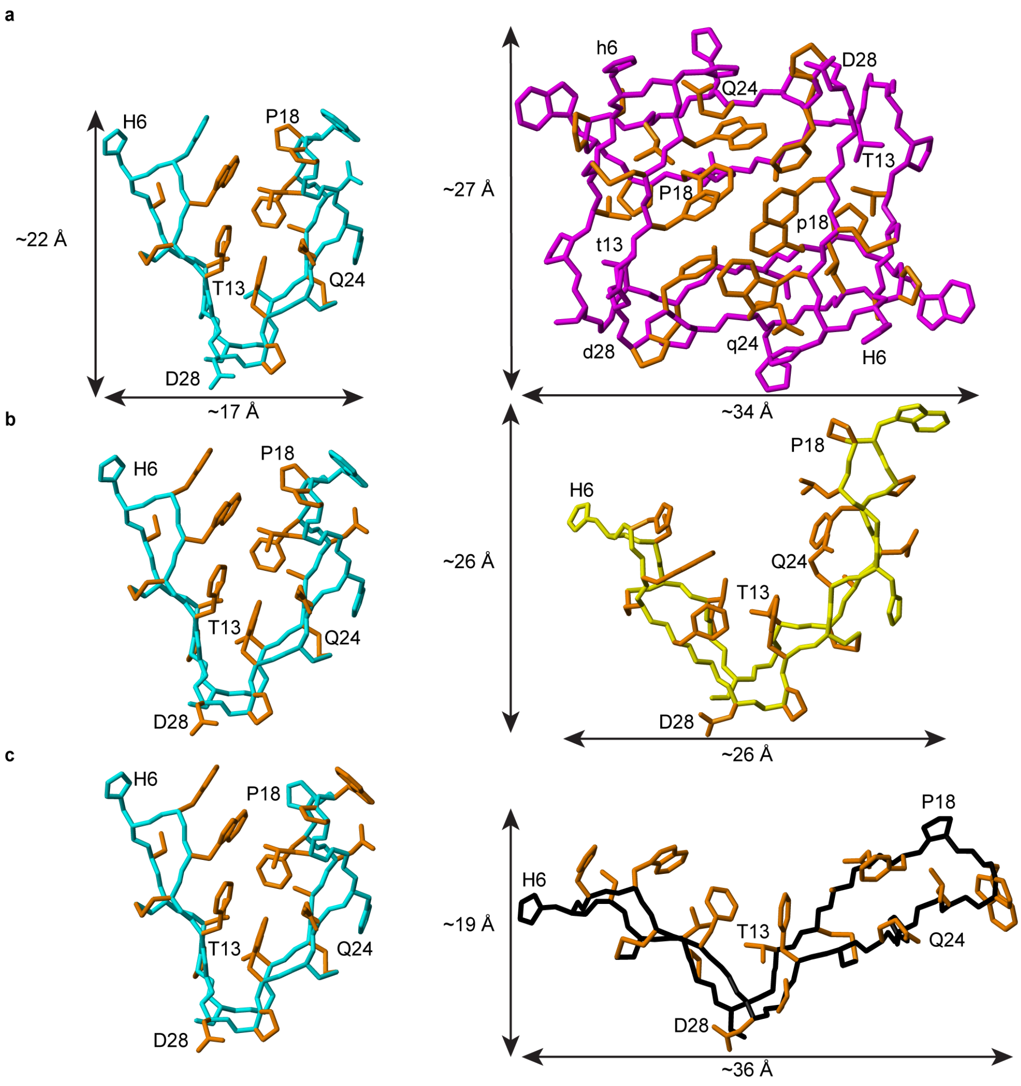


**Supplementary Figure 7: Three-dimensional structures of PDP-23 in different environments. (a)** Comparison of key structural differences between monomeric PDP-23 when in 80:20 $H_2O/CD_3CN$ and dimeric PDP-23 when in 90:10 $H_2O/D_2O$. PDP-23 in (80:20) $H_2O/CD_3CN$ is colored cyan, PDP-23 in 90:10 $H_2O/D_2O$ is colored magenta. Sidechains of residues showing chemical shift differences >0.15 ppm between the two conditions are highlighted in orange in both structures, reflecting changes in the core. **(b)** Comparison of key structural differences between PDP-23 when in (80:20) $H_2O/CD_3CN$ and when exposed to SDS micelles. PDP-23 in 80:20 $H_2O/CD_3CN$ is colored cyan, PDP-23 exposed to SDS micelles is colored yellow. Residues with chemical shift deviations >0.15 ppm are highlighted in orange in both structures. **(c)** Comparison of key structural differences between PDP-23 when in 80:20 $H_2O/CD_3CN$ and when exposed to DPC micelles. PDP-23 in 80:20 $H_2O/CD_3CN$ is colored cyan, PDP-23 exposed to DPC micelles is colored black. Residues with chemical shift deviations >0.15 ppm are highlighted in orange in both structures. For all structures residue labels are supplied for orientation in the case of PDP-23 in 90:10 $H_2O/D_2O$ capital and non-capital letters are used to denote individual monomers.

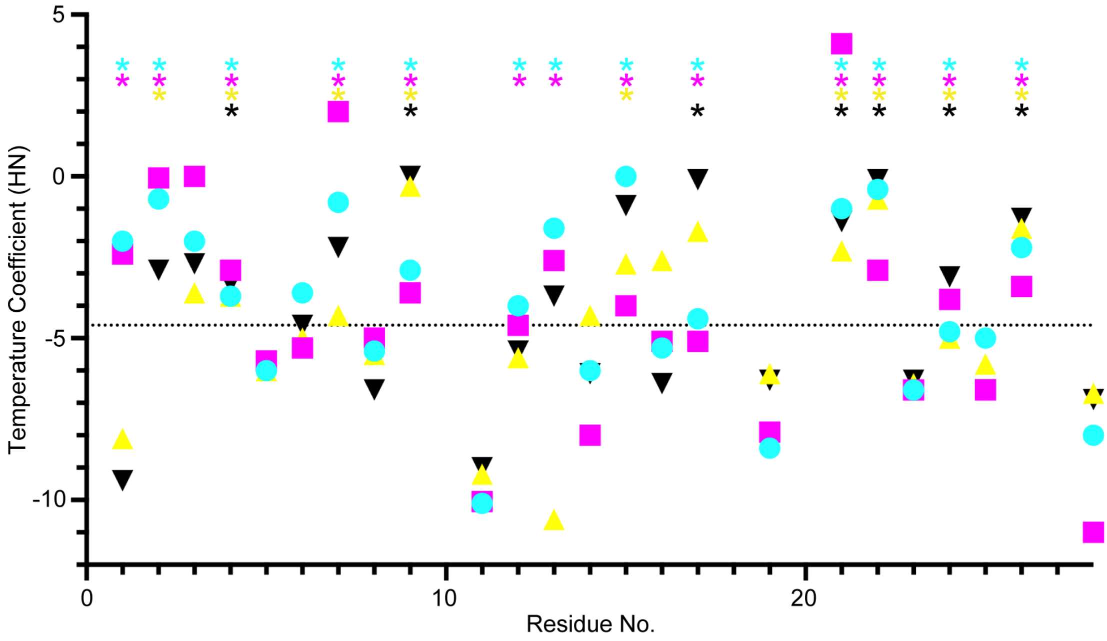


**Supplementary Figure 8: Amide proton temperature coefficients for PDP-23 in different environments.** Amide protons involved in hydrogen bonds are protected from solvent and their chemical shift is less affected by temperature changes. For amide protons with a temperature coefficient >-4.6 ppb/K (indicated by the dashed line) the likelihood of being involved in a hydrogen bond is >86%[1]. Data points are color coded according to conditions; 80:20 $H_2O/CD_3CN$ (cyan), 90:10 ($H_2O/D_2O$) (magenta), SDS micelles (yellow), DPC micelles (black). Hydrogen bonds with confirmed unambiguous acceptors are highlighted by * at the top, and in all cases, these are consistent between conditions. Based on temperature coefficient analysis the majority of hydrogen bonds are unaffected by condition changes, but residues like Gly1 and Thr13 do show a larger change in temperature dependence indicating hydrogen bonds are being broken in micelles.

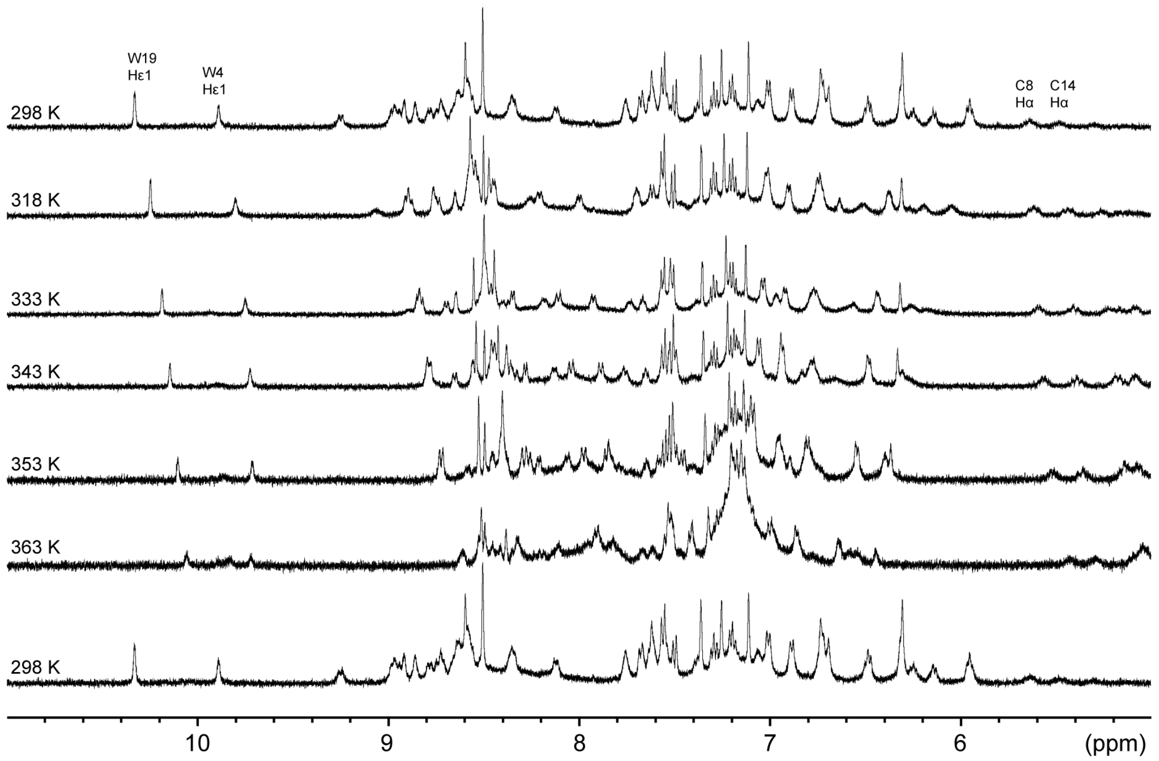


**Supplementary Figure 9: One-dimensional $^1$H NMR spectra showing thermal stability of PDP-23.** 1D $^1$H NMR spectra of PDP-23 recorded while cycling the temperature from 298-363-298 K. Data show only partial denaturing effects at a temperature of 363 K at the two tryptophan indole signals and Cys Hα signals. In contrast, aromatic signals around 5.8-6.5 ppm, which are generated from the packing of Phe sidechains in the dimer core, move downfield, consistent with the effect of addition of acetonitrile and separation into monomers. The complete reversibility of thermal denaturation was confirmed when the temperature was again decreased to 298 K, with the 1D spectrum being identical to the 1D spectrum prior to heating.

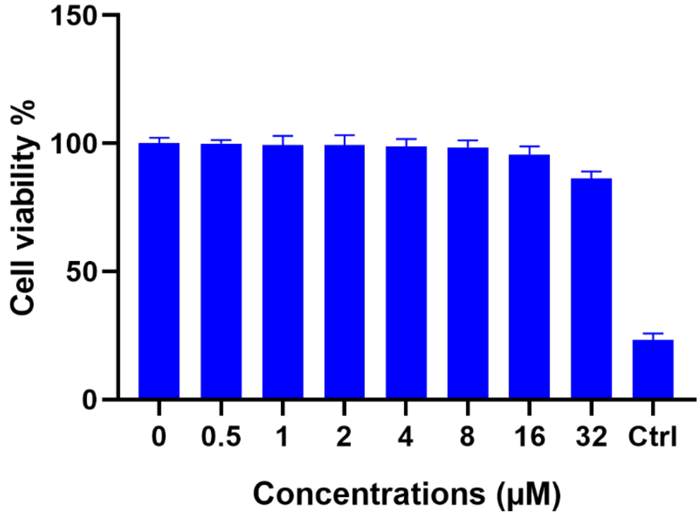


**Supplementary Figure 10: Cell viability of HeLa cells exposed to PDP-23 for 48 hours.** Cells were incubated in a serial dilution of PDP-23 for 48 h and cell viability was then determined by MTT assay. Cells treated with 5% (v/v) DMSO were used as a positive control. The assay was performed in triplicate and the results are presented as the means ± SD.

**Supplementary Table 1: Structural Statistics of PDP-23 in different environments**

| | 80:20 $H_2O/CD_3CN$ | 90:10 $H_2O/D_2O$ | SDS micelles | DPC micelles |
|---|---|---|---|---|
| **Energies (kcal/mol)** | | | | |
| Overall | -801.1 ± 12.0 | -1743.1 ± 36.0 | -789.7 ± 21.4 | -770.7 ± 37.5 |
| Bonds | 12.60 ± 0.98 | 20.97 ± 1.17 | 11.74 ± 1.27 | 11.22 ± 1.02 |
| Angles | 38.26 ± 2.07 | 54.07 ± 4.78 | 31.37 ± 4.13 | 32.45 ± 8.90 |
| Improper | 13.36 ± 1.59 | 21.86 ± 2.85 | 14.56 ± 3.87 | 12.99 ± 4.09 |
| Dihedral | 121.8 ± 1.38 | 238.8 ± 1.63 | 119.1 ±1.26 | 119.8 ± 2.26 |
| Van der Waals | -96.51 ± 3.02 | -272.3 ± 8.71 | -97.43 ± 7.75 | -98.59 ± 10.9 |
| Electrostatic | -891.1 ± 10.8 | -1806.8 ± 30.3 | -869.5 ± 19.5 | -848.9 ± 32.8 |
| NOE | 0.16 ± 0.02 | 0.07 ± 0.06 | 0.36 ± 0.97 | 0.06 ± 0.05 |
| Cdih | 0.42 ± 0.21 | 0.22 ± 0.15 | 0.18 ± 0.57 | 0.17 ± 0.12 |
| **MolProbity Statistics** | | | | |
| Clashes (>0.4 Å/1000 atoms) | 15.7 ± 4.6 | 11.5 ± 2.5 | 12.8 ± 3.8 | 12.6 ± 4.4 |
| Poor Rotamers | 0 | 0.3 ± 0.5 | 0 | 0.5 ± 0.6 |
| Ramachandran Outliers (%) | 0 | 0 | 1.2 ± 1.8 | 2.9 ± 3.3 |
| Ramachandran Favored (%) | 97.7 ± 1.9 | 97.7 ± 1.8 | 93.7 ± 5.2 | 81.2 ± 5.6 |
| MolProbity Score | 1.84 ± 0.2 | 1.64 ± 0.1 | 1.94 ± 0.3 | 2.52 ± 0.4 |
| MolProbity score percentile | 82.7 ± 8.5 | 91.0 ± 3.8 | 77.0 ± 13.3 | 47.4 ± 18.6 |
| **Atomic RMSD (Å)** | | | | |
| Mean Global Backbone | 0.43 ± 0.11 | 0.73 ± 0.15 | 1.60 ± 0.50 | 1.82 ± 0.63 |
| Mean Global Heavy | 0.92 ± 0.15 | 1.28 ± 0.15 | 2.66 ± 0.61 | 2.78 ± 0.82 |
| **Experimental Restraints** | | | | |
| **Distance Restraints** | | | | |
| Short range (i-j < 2) | 217 | 436 | 211 | 192 |
| Medium range (i-j < 5) | 74 | 114 | 39 | 44 |
| Long range (i-j > 5) | 115 | 192 | 77 | 59 |
| Intermolecular | | 162 | | |
| Hydrogen bonds | 26 (13 H-bonds) | 52 (26 H-bonds) | 20 (10 H-bonds) | 12 (6 H-bonds) |
| Total | 432 | 956 | | 307 |
| **Dihedral Restraints** | | | | |
| Φ | 18 | 40 | 3 | 11 |
| Ψ | 20 | 44 | 2 | 11 |
| Total | 38 | 84 | 5 | 22 |
| **Restraint violations** | | | | |
| Total NOE violations > 0.2 Å | 0 | 0 | 0 | 0 |
| Total Dihedral violations > 2.0° | 0 | 0 | 0 | 0 |

**Supplementary Table 2: Chemical Shift Differences between 90:10 ($H_2O/D_2O$) and 80:20 ($H_2O/CD_3CN$)**

| Residue | Atom | Δδ ($H_2O/D_2O$)-($H_2O/CD_3CN$) (>0.15 ppm are listed) |
|---|---|---|
| **1** | HA3 | -0.178 |
| **2** | HB2 | 0.284 |
| **2** | QD | 0.248 |
| **2** | HN | 0.166 |
| **3** | HB3 | 0.206 |
| **4** | HD1 | 0.412 |
| **4** | HB2 | 0.259 |
| **4** | HB3 | 0.259 |
| **4** | HZ3 | -0.29 |
| **4** | HZ2 | -0.373 |
| **4** | HH2 | -0.448 |
| **4** | CB | 1.559 |
| **6** | HA | 0.16 |
| **6** | HN | -0.226 |
| **7** | HB3 | 0.265 |
| **7** | HB2 | 0.254 |
| **7** | HN | 0.2 |
| **9** | HB | -0.273 |
| **11** | HN | -0.287 |
| **12** | HN | -0.176 |
| **14** | HB3 | 0.333 |
| **14** | HN | -0.265 |
| **14** | CB | 1.131 |
| **15** | QB | 0.595 |
| **16** | CB | 2.376 |
| **16** | CA | 1.446 |
| **17** | HZ | 0.709 |
| **17** | QE | 0.614 |
| **17** | QD | 0.257 |
| **17** | HB2 | -0.193 |
| **17** | HN | -0.244 |
| **17** | CB | 1.383 |
| **18** | HG2 | -0.156 |
| **18** | HD2 | -0.162 |
| **18** | HA | -0.207 |
| **20** | HB3 | -0.228 |
| **21** | HN | 0.525 |
| **21** | HB2 | 0.293 |
| **21** | HG | 0.173 |
| **21** | HB3 | 0.153 |
| **22** | HN | -0.294 |
| **24** | HE21 | 0.289 |
| **24** | HE22 | 0.193 |
| **24** | HG3 | -0.355 |
| **25** | HN | 0.263 |
| **26** | QD | -0.151 |
| **26** | HZ | -0.373 |
| **26** | QE | -0.714 |
| **27** | HD3 | 0.325 |
| **27** | HD2 | 0.308 |
| **27** | HB3 | 0.16 |
| **27** | CA | 1.947 |
| **27** | CB | 1.527 |
| **28** | HN | 0.414 |
| **28** | CB | 1.823 |
| **28** | CA | 1.092 |

**Supplementary Table 3: Chemical shift difference 90:10 ($H_2O/D_2O$) and SDS/DPC micelles. * denotes the difference being <0.15 ppm**

| Residue | Atom | Δδ ($H_2O/D_2O$)-($H_2O$/SDS), ($H_2O/D_2O$)-($H_2O$/DPC) (>0.15 ppm are listed) |
|---|---|---|
| 1 | HA3 | 0.31, 0.37 |
| 1 | HA2 | *, -0.22 |
| 1 | HN | -0.38, -0.58 |
| 2 | HA | -0.19, * |
| 2 | HN | *, -0.27 |
| 2 | HE* | -0.279, -0.453 |
| 2 | HD* | -0.638, -0.735 |
| 2 | HB2 | -0.82, -0.8 |
| 2 | HB3 | -1.25, -1.134 |
| 2 | CB | 1.154, 3.949 |
| 3 | HA | -0.219 -0.302 |
| 3 | HB3 | -0.298, -0.215 |
| 3 | HB2 | -0.332, 0.295 |
| 3 | HN | -0.562, -0.656 |
| 4 | HH2 | 0.536, 0.583 |
| 4 | HZ3 | 0.351, 0.394 |
| 4 | HZ2 | 0.301, 0.273 |
| 4 | HE3 | 0.171, 0.199 |
| 4 | HB3 | -0.366, -0.291 |
| 4 | HB2 | -0.419, -0.384 |
| 4 | HD2 | -0.575, -0.547 |
| 4 | HE1 | *, -0.539 |
| 4 | HN | *, -0.242 |
| 4 | CB | 1.709, 4.504 |
| 5 | HD2 | 0.338, 0.421 |
| 5 | HB3 | 0.216, * |
| 5 | HB2 | 0.173, * |
| 5 | HN | *, -0.311 |
| 5 | CA | *, 3.006 |
| 5 | CB | *, 1.986 |
| 6 | HN | 0.195, * |
| 6 | HA | -0.538, -0.263 |
| 6 | CA | *, 1.765 |
| 6 | CB | -1.03, * |
| 7 | HB3 | -0.336, -0.331 |
| 7 | HB2 | -0.36, -0.412 |
| 7 | HN | -0.52, -0.498 |
| 7 | CA | *, 2.522 |
| 8 | HA | 0.258, 0.163 |
| 8 | HN | *, -0.162 |
| 8 | CA | *, -5.809 |
| 8 | CB | -8.604, * |
| 9 | HB | 0.957, 0.774 |
| 9 | HA | 0.593, 0.577 |
| 9 | HG2* | 0.308, 0.284 |
| 9 | HG1* | 0.254, 0.238 |
| 9 | HN | 0.179, 0.162 |
| 10 | HA | 0.214, 0.162 |
| 10 | HG3 | 0.202, 0.178 |
| 11 | CA | *, 2.89 |
| 12 | HA2 | 0.272, * |
| 13 | HG* | 0.168, 0.191 |
| 13 | HN | -0.291, -0.966 |
| 13 | CB | *, 2.6 |
| 14 | HN | 0.2, * |

| | | |
|---|---|---|
| 14 | HB3 | -0.44, -0.418 |
| 15 | HB* | -0.555, -0.498 |
| 15 | CA | *, 2.613 |
| 15 | CB | 7.354, * |
| 16 | HA | 0.205, 0.208 |
| 16 | HB3 | -0.29, -0.28 |
| 16 | HB2 | -0.291, -0.28 |
| 16 | HN | *, -0.677 |
| 16 | CB | 2.912, 5.707 |
| 17 | HN | 0.718, 0.728 |
| 17 | HB2 | *, -0.191 |
| 17 | HD* | -0.301, -0.299 |
| 17 | HE* | -0.689, 0.663 |
| 17 | HZ | -0.879, -0.824 |
| 17 | CA | *, 5.004 |
| 17 | CB | 2.209, * |
| 18 | HG3 | 0.182, * |
| 19 | HZ3 | *, 0.244 |
| 19 | HH2 | *, 0.167 |
| 19 | HE1 | *, -0.598 |
| 19 | CA | -1.913, * |
| 20 | HD2 | 1.851, 0.25 |
| 20 | HD3 | 1.69, * |
| 20 | HG2 | 0.416, 0.188 |
| 20 | HG3 | *, 0.514 |
| 20 | HB3 | 0.309, 0.646 |
| 20 | HA | *, 0.187 |
| 20 | CA | *, -9.006 |
| 21 | HG | *, 0.156 |
| 21 | HB3 | -0.288, -0.264 |
| 21 | HB2 | -0.324, -0.326 |
| 21 | HN | -0.532, -0.618 |
| 21 | CA | -2.427, * |
| 22 | HN | *, 0.178 |
| 22 | HA3 | -0.177, -0.167 |
| 23 | HN | 0.16, * |
| 23 | CB | *, 2.731 |
| 24 | HE21 | 0.222, 0.236 |
| 24 | HE22 | 0.181, * |
| 24 | HA | -0.239, -0.228 |
| 24 | HB3 | -0.252, -0.408 |
| 24 | HB2 | -0.419, -0.238 |
| 24 | HG3 | -0.482, -0.411 |
| 24 | HG2 | -0.452, -0.461 |
| 24 | CA | -2.222, * |
| 25 | HA | -0.342, -0.338 |
| 25 | CA | *, 1.865 |
| 26 | HA | 0.159, 0.324 |
| 26 | HB3 | -0.178, -0.174 |
| 26 | HB2 | -0.232, * |
| 26 | HD* | -0.368, -0.306 |
| 26 | HZ | -0.919, -0.835 |
| 26 | HE* | -1.14, -1.09 |
| 27 | HD2 | 0.325, 0.495 |
| 27 | HD3 | 0.257, 0.281 |
| 27 | CA | *, 3.628 |
| 28 | HN | 0.597, 0.496 |
| 28 | HA | 0.34, 0.275 |
| 28 | HB3 | -0.31, * |
| 28 | CB | -8.252, -5.457 |

**Supplementary Reference**

1. Cierpicki, T. & Otlewski, J. Proton temperature coefficients as hydrogen bond indicators in proteins. J. Biomol. NMR. **21**, 249-261, (2001).
